# Identification and characterization of a new soybean promoter induced by *Phakopsora pachyrhizi*, the causal agent of Asian soybean rust

**DOI:** 10.1101/2020.06.07.138586

**Authors:** Lisa Cabre, Stephane Peyrard, Catherine Sirven, Laurine Gilles, Bernard Pelissier, Sophie Ducerf, Nathalie Poussereau

## Abstract

**Background:** *Phakopsora pachyrhizi* is a biotrophic fungal pathogen responsible for the Asian soybean rust disease causing important yield losses in tropical and subtropical soybean-producing countries. *P. pachyrhizi* triggers important transcriptional changes in soybean plants during infection, with several hundreds of genes being either up- or downregulated.

**Results:** Based on published transcriptomic data, we identified a predicted chitinase gene, referred to as *GmCHIT1*, that was upregulated in the first hours of infection. We first confirmed this early induction and showed that this gene was expressed as early as 8 hours after *P. pachyrhi*zi inoculation. To investigate the promoter of *GmCHIT1*, transgenic soybean plants expressing the green fluorescence protein (GFP) under the control of the *GmCHIT1* promoter were generated. Following inoculation of these transgenic plants with *P. pachyrhizi*, GFP fluorescence was detected in a limited area located around appressoria, the fungal penetration structures. Fluorescence was also observed after mechanical wounding whereas no variation in fluorescence of p*GmCHIT1*:GFP transgenic plants was detected after a treatment with an ethylene precursor or a methyl jasmonate analogue.

**Conclusion:** We identified a soybean chitinase promoter exhibiting an early induction by *P. pachyrhizi* located in the first infected soybean leaf cells. Our results on the induction of *GmCHIT1* promoter by *P. pachyrhizi* contribute to the identification of a new pathogen inducible promoter in soybean and beyond to the development of a strategy for the Asian soybean rust disease control using biotechnological approaches.

## BACKGROUND

Rusts are among the most damaging crop diseases, causing very severe losses in crop yield ^1^. In particular, Asian soybean rust is the most destructive foliar disease of soybean (*Glycine max* (L.) Merr.) and is caused by the biotrophic basidiomycete fungus *Phakopsora pachyrhizi* Syd. & P. Syd ^2^. Initially localized in Asia, *P. pachyrhizi* has spread across the world and reached the South American continent in the 2000s, bringing important economic losses to soybean growers. Brazil, one of the leading soybean-producing countries, is impacted by the disease each year. Highest damages on grain harvest between 2007 and 2014 reached 571.8 thousand tons, e.g., 6% of the national production ^3^.Infection by *P. pachyrhizi* starts with the germination of uredospores on the soybean leaf, leading to the formation of an appressorium. From the appressorium *P. pachyrhizi* penetrates directly into the epidermal cells of its hosts. Between 24 and 48 hours later, fungal hyphae colonized infected tissues and haustoria are observed in the mesophyll cells. Approximately 5-8 days post infection, uredinia appear on the abaxial side of the leaves and new urediniospores are released, leading to inoculation of healthy plants through airborne spore dissemination ^4^. Symptoms are characterized by tan-coloured lesions and chlorosis of the leaves. In the most severe cases, defoliation and quick maturation of soybean with a reduction of seed size and weight can be observed in a few days after initial infection ^5,6^.

Today, the control of *P. pachyrhizi* is essentially based on fungicidal treatments. Demethylation inhibitors (DMIs) impairing sterol biosynthesis, as well as succinate dehydrogenase inhibitors (SDHIs) and quinone outside inhibitors (QoIs), blocking mitochondrial respiration, are the most commonly used fungicides ^7,8^. However, the repetitive use of molecules with these three modes of action and the fungicide adaptation capability of the pathogen have resulted in a decrease in treatment efficacy ^3^. Genetic resistance of soybean to *P. pachyrhizi* is well documented and could be seen as an alternative to the use of pesticides. Thus far, seven dominant R genes, named *Rpp1* to *Rpp7*, have been identified ^9–13^. However, these resistance genes are only effective against specific isolates of *P. pachyrhizi* ^14^ and the resistance conferred by these genes can be easily overcome, making breeding solutions very challenging ^15^. Today, no soybean cultivars resistant to most of the rust isolates are available. In this context, biotechnological approaches are foreseen as alternative solutions to control Asian soybean rust ^8,16^.

A common strategy in plant engineering for disease resistance is to overexpress a defence-related gene placed under the control of a constitutive promoter. However, permanent and high ectopic expression of such a gene can impact the plant’s fitness and development ^17^. These challenges can be overcome by using a pathogen-inducible promoter allowing transgene expression only when and where it is needed. The advantage of these regulated promoters is well illustrated by the expression of the multi-pathogen resistant gene *Lr34res* in barley ^18^. The *Lr34res* gene encoding an ATP-binding cassette (ABC) transporter was originally identified in wheat as providing durable resistance to 3 wheat rusts (*Pucccinia triticina, P.striiformis, P.graminis*) and the powdery mildew (*Blumeria graminis* f.sp. *tritici*). This gene was successfully transferred in barley and conferred resistance against *Puccinia hordei* and the powdery mildew *Blumeria graminis f.sp. hordei*. However, *Lr34res* expression controlled by its native promoter resulted in negative effect on plant growth and fitness ^19^. To avoid these pleiotropic effects, Boni *et al*. (2018) developed transgenic barley expressing the *Lrs34 res* gene placed under the control of the barley germin-like GER4 promoter, a pathogen inducible promoter ^20^. They observed that the negative pleiotropic effects were reduced compared to barley plants containing the same gene placed under control of its native promoter. The composition of the pathogen-inducible promoters has also to be considered since the promoter region may contain several *cis*-regulatory elements such as binding sites for transcription factors and/or regulatory proteins. These elements that regulate gene expression patterns can be activated by different stimuli ^21^. As a consequence, pathogen-inducible promoters are often induced by other stimuli such as wounding and/or hormones. Many pathogen-inducible promoters have been studied in different plants ^20,22,23^, but very few have been reported in soybean. For instance, *GmPPO12* (*Glyma04g14361*) promoter controlling a polyphenol oxidase has been found to be rapidly and strongly induced by *Phytophthora sojae* in transformed soybean hairy roots and two regions were identified as essential for promoter activity ^24^. In addition, Liu *et al.* (2014) discovered 23 *cis*-regulatory elements responsible for the induction of several genes by the soybean cyst-nematode *Heterodera glycines* ^25^ and they proposed to consider them for synthetic promoter engineering.

Plant responses to pathogen attacks involve the activation of a set of genes coding for different proteins. Among them, pathogenesis-related (PR) proteins are produced and highly accumulated ^26^. Chitinases represent a subset of pathogenesis-related proteins. These enzymes that belong to families 18 and 19 of the glycosyl hydrolases^27^, have the ability to randomly hydrolyse beta-1,4-glycoside bonds of chitin, a major component of the fungal cell wall. The resulting chitin fragments act as a potent pathogen-associated molecular pattern (PAMP) that induces PAMP-triggered immunity ^27^. Plant chitinases have also been shown to be implicated in the defence against insects; in response to abiotic stresses such as cold, drought or metal toxicity; and in plant development ^28,29^.

In this publication, we report the identification and characterization of the soybean chitinase promoter p*GmCHIT1* that we selected from a set of transcriptomic data^30,31^. This promoter drives both early and late overexpression of a chitinase encoding gene upon *P. pachyrhizi* infection. Its specificity to fungal exposure versus activation by different hormonal and abiotic stress pathways was evaluated through the generation of stable transgenic soybeans harbouring a p*GmCHIT1*:GFP fusion. Our study was carried out on the *P. pachyrhizi* / soybean pathosystem, allowing induction of the promoter by the pathogen in the crop of interest. To our knowledge, this is the first characterization of a soybean promoter inducible by Asian soybean rust.

## RESULTS

### The soybean chitinase gene *GmCHIT1* is induced by Asian soybean rust

Several transcriptomic data on soybean gene expression during *P. pachyrhizi* infection have been generated and published. In 2010, Tremblay *et al.* used DNA array to analyse gene expression in the palisade and mesophyll cells infected by the pathogen. They identified 685 upregulated genes 10 days after soybean rust inoculation (dpi), and most of them were related to plant defence response and metabolism^30^. In 2011, they used next-generation sequencing (NGS) to analyse soybean gene expression patterns in leaves and described 1,713 genes upregulated 10 dpi, with many of them encoding proteins involved in metabolism and transport ^31^. Considering that upregulated genes are a potential source of inducible promoters, we searched for genes upregulated in both experiments. We identified 220 common upregulated genes, and a ranking of these genes according to their fold change was determined for each experiment (see additional file 1: Table S1). Among the commonly upregulated genes, one-quarter (26%) were associated with metabolism function, 18% were implicated in signal transduction and 12% were annotated as transporters (see additional file 2: Figure S1). Eleven plant defence-related genes representing 5% of the commonly upregulated genes were also identified. Among them, two genes annotated as predicted chitinase (*Glyma.13G346700* and *Glyma.11G124500*) were highly induced at 10 dpi. They were also described as up-regulated 24 h post-infection, in agreement with SoyKB data (http://soykb.org/). Moreover, according to internal transcriptomic data, *Glyma.11G124500* revealed no induction after treatment with a chitin oligosaccharide (the chitin heptaose) unlike *Glyma.13G346700* (see additional file 3: Figure S2). Chitin is a major component of the fungal cell wall and can be detected by the host plant as a PAMP. Therefore, we selected *Glyma.11G124500* as potentially specifically induced by *P. pachyrhizi* during early (24 h) and late stages of infection (10 days).

*Glyma.11G124500*, located on chromosome 11, includes a coding sequence of 705 bp with two exons, a 5’UTR of 57 bp and a 3’UTR of 217 bp. This gene encodes a protein (Glyma.11G124500 1. p) of 235 amino acids with a glycosyl hydrolase motif of family 19 (PF00182 domain from amino acid 38 to 235) and was annotated as a chitinase. This putative function was reinforced by a sequence comparison (see additional file 4: Figure S3 and additional file 5: Figure S4). *Glyma.11G124500* was therefore renamed *GmCHIT1*.

Expression of *GmCHIT1* during infection of wild type soybean leaves by *P. pachyrhizi* was then monitored by RT-qPCR. *GmCHIT1* was expressed as early as 8 hpi (hours post-inoculation) (2.5-fold compared to the mock treatment), and its expression increased during infection reaching 6-7-fold compared to the mock treatment at 1-3 dpi. The highest level of *GmCHIT1* expression (300-fold compared to healthy leaves) was observed at a late stage of infection when the inoculated leaves were totally chlorotic and covered with sporulating uredinia (10 dpi) (Figure 1a). In our conditions, no visual symptoms were observed at 8 hpi and uredia appeared at 6/7 dpi, revealing that the gene was induced before the emergence of disease symptoms (Figure 1a, b). According to the expression results, we selected the *GmCHIT1* promoter as a good candidate induced by *P. pachyrhizi.*

**Figure 1:**
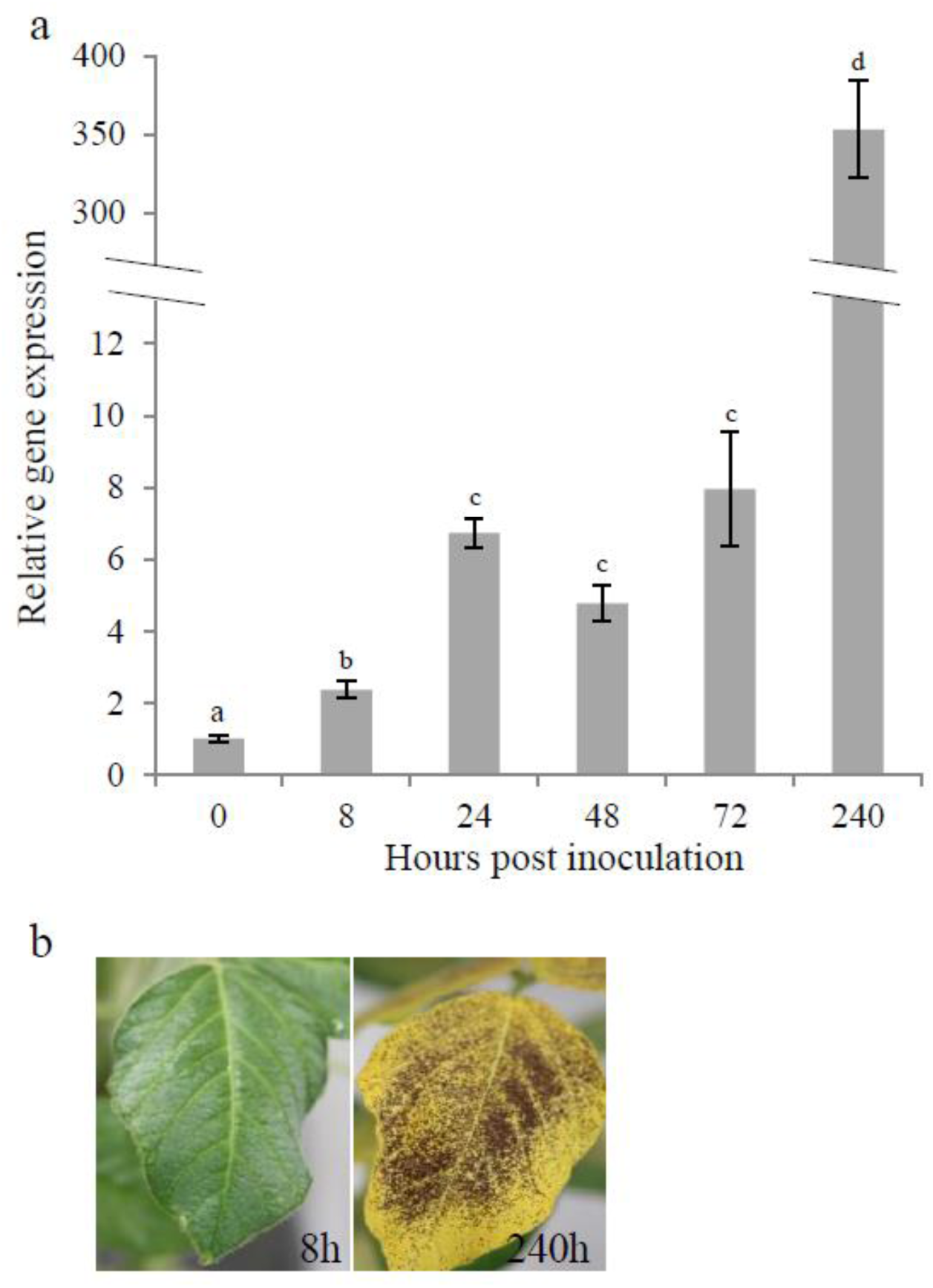
Relative expression of *GmCHIT1* in soybean leaves during *P. pachyrhizi* infection. (a) Quantitative RT-qPCR analysis was carried out to quantify *GmCHIT1* transcript accumulation at 0, 8, 24, 48, 72 and 240 hpi compared to that in the mock-treated plants. The actin (GenBank: NM_001289231.2) and an unknown protein (GenBank: BE330043) ^68^ encoding genes were used as references. Three independent biological replicates ± standard errors. Different letters indicate a significant difference determined by a Student’s t-test (*p* < 0.05) between time point of inoculation. (b) Stages of infection of soybean leaves at 8 and 240 hpi.

### Analysis of the activity of the *GmCHIT1* promoter in response to *P. pachyrhizi* inoculation

To study the expression and inducibility of *GmCHIT1* promoter, a fragment of 3454 bp upstream of the coding sequence was selected. Indeed, analysis of this sequence with PLACE software ^32^ revealed several cis-regulatory elements related to pathogen infection (see additional file 6: Figure S5). Twenty-one W boxes (TGAC) ^33^ and 10 GT1 boxes (GAAAAA) ^34^ were identified. Five MYB recognition elements (GGATA)^35^ were also found as well as two auxin (TGTCTC and KGTCCCAT) ^36^ and two gibberellic acid-responsive elements (CAACT) ^37^.

The activity of the *GmCHIT1* promoter following *P. pachyrhizi* inoculation was then evaluated via the generation of reporter stable transgenic soybeans. For this, the selected promoter region was fused to the GFP reporter gene (p*GmCHIT1*:GFP), and transgenic plants were selected. *P. pachyrhizi* spores were sprayed on the plants, and fluorescence surrounding the infection spots was clearly observed at 24 and 72 hpi in three independent p*GmCHIT1*:GFP lines (Figure 2a (line 131); additional file 7 Figure S6 (lines 129 and 133)). However, a low GFP signal was also observed in leaf veins in the absence of the fungal infection, revealing a basal expression of the promoter in fully developed 3-week-old soybean plants. *GFP* expression was followed by RT-qPCR and a low induction was detected at 72 hpi (Figure 2b). Western blot analysis revealed the presence of GFP in non-infected leaves and an accumulation of GFP-protein at 72 hpi (Figure 2c). To precise this over-accumulation, a confocal microscopy study was conducted on line 131 at 24 hpi when spores have germinated and differentiated appressoria. GFP fluorescence was particularly detectable around the fungal pathogen and more precisely in cells located around appressoria, the fungal penetration structures (Figure 3).

**Figure 2:**
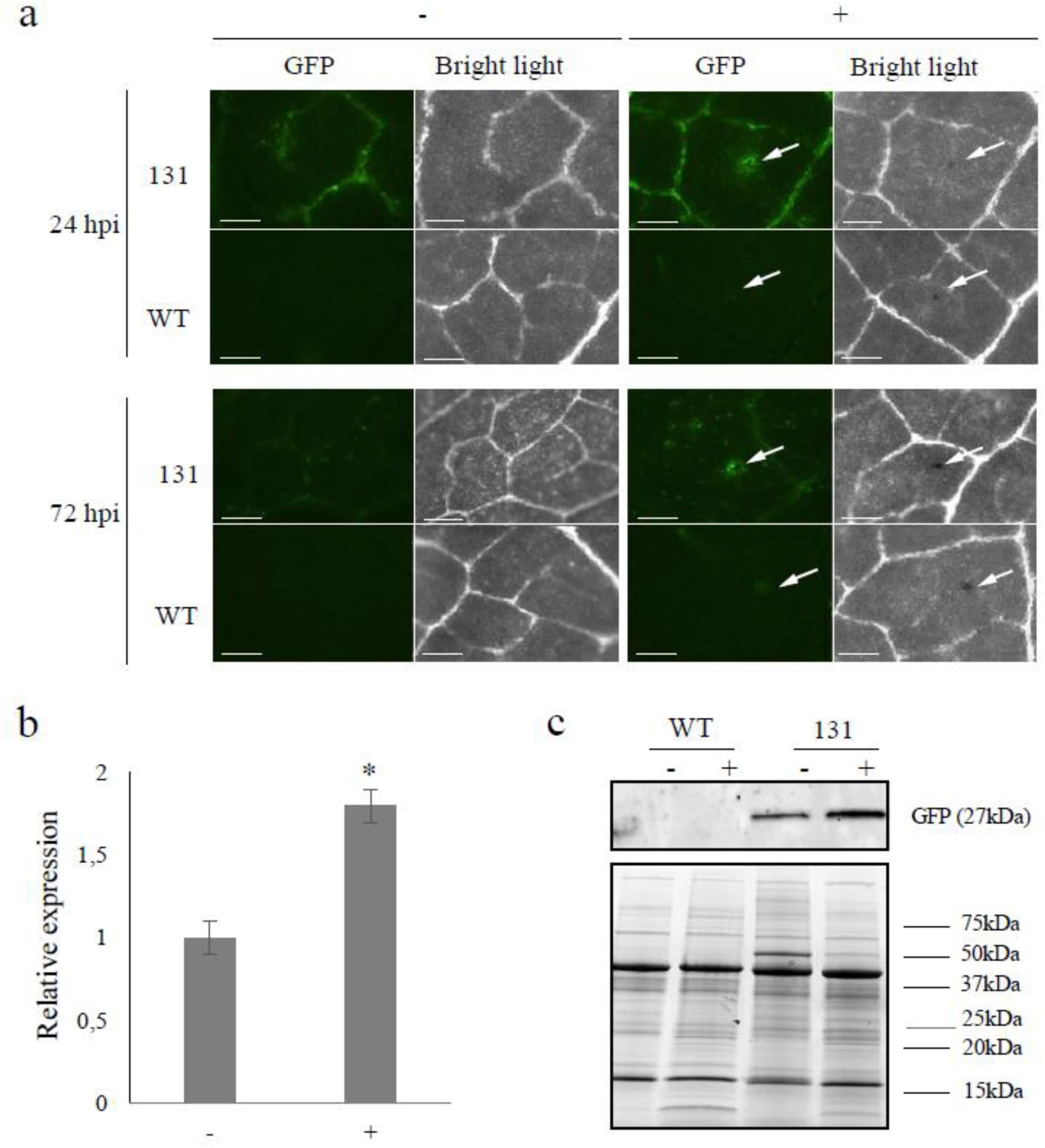
Detection of GFP in stable transgenic soybeans. (a) Leaves of T1 lines 131 transformed with the p*GmCHIT1*:GFP construction were observed using a dissection scope (Leica Z16 APO) under GFP filter and bright light at 24 and 72 hours after. *P. pachyrhizi* inoculation (+) or mock treatment (-). Arrows indicate the inoculation spots. Bar-scales represent 200 µm. (b) Relative expression of *GFP* in 131 line (*pGmCHIT:GFP*) at 72 h after. *P. pachyrhizi* inoculation (+) and non-infected (-) plants. The actin (GenBank: NM_001289231.2) and an unknown protein (GenBank: BE330043) ^68^ encoding genes were used as references. Three independent biological replicates ± standard errors are shown. *: significant difference between treated (+) and untreated (-) leaves determined by a Student’s t-test (*p* < 0.05). (c) Detection of GFP protein 72 hours after. *P. pachyrhizi* inoculation in WT and 131 line plants by immunoblotting with an antibody raised against the GFP. Homogenous loading was checked on the gel by Strain Free detection technology (Biorad, US) (below).

**Figure 3:**
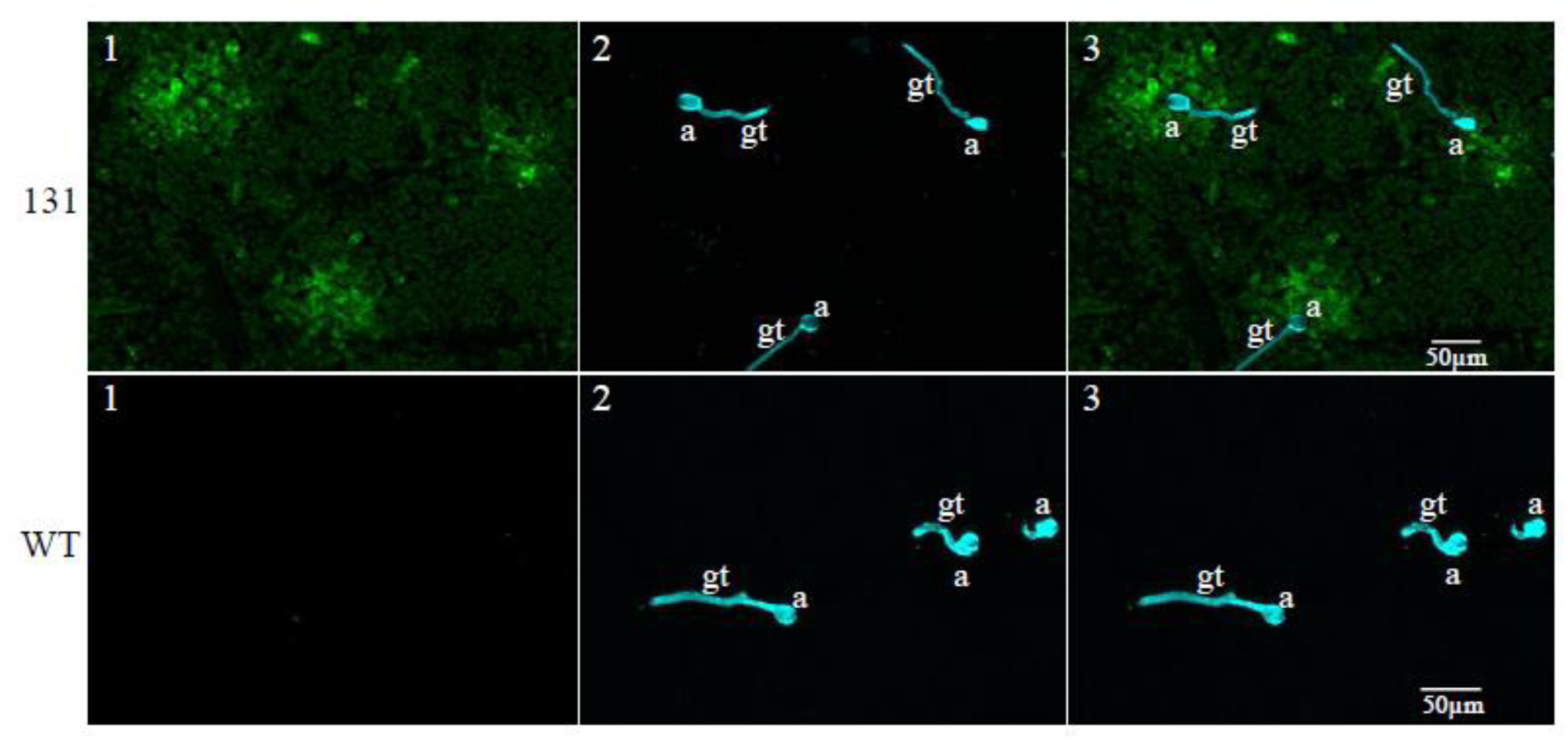
Representative confocal image (z-stack projection) showing GFP induction around *P. pachyrhizi* appressoria at 24 hpi. Fungal structures on the leaf surface are stained in blue with calcofluor. a: appressoria, gt: germ tube. Picture 1: GFP detection. Picture 2: calcofluor staining. Picture 3: merging of pictures 1 and 2. The observations were conducted on 131 (p*GmCHIT1*:GFP) and WT plants.

### Activity of the *GmCHIT1* promoter in different soybean tissues

To determine the tissue specificity of the chitinase promoter, GFP fluorescence of plants from the 131 line was investigated in roots, young leaves and flowers of non-infected plants. p*CsVMV*:GFP plants containing the strong constitutive Cassava Vein Mosaic Virus promoter were used as a positive control. As expected, a strong GFP fluorescence was observed in all analysed tissues of plants transformed with p*CsVMV*:GFP, whereas no GFP signal was detected in WT soybean plants (Figure 4). In the case of plants transformed with p*GmCHIT1*:GFP (line 131), a light GFP signal was detected in primary and some lateral roots. While GFP expression was observed in veins of developed leaves (Figures 2, 5 and 6), no signal was detectable in young leaves at this magnification (Figure 4). This low detection of GFP could be considered as the baseline expression of the *GmCHIT1* promoter in the different tissues observed.

**Figure 4:**
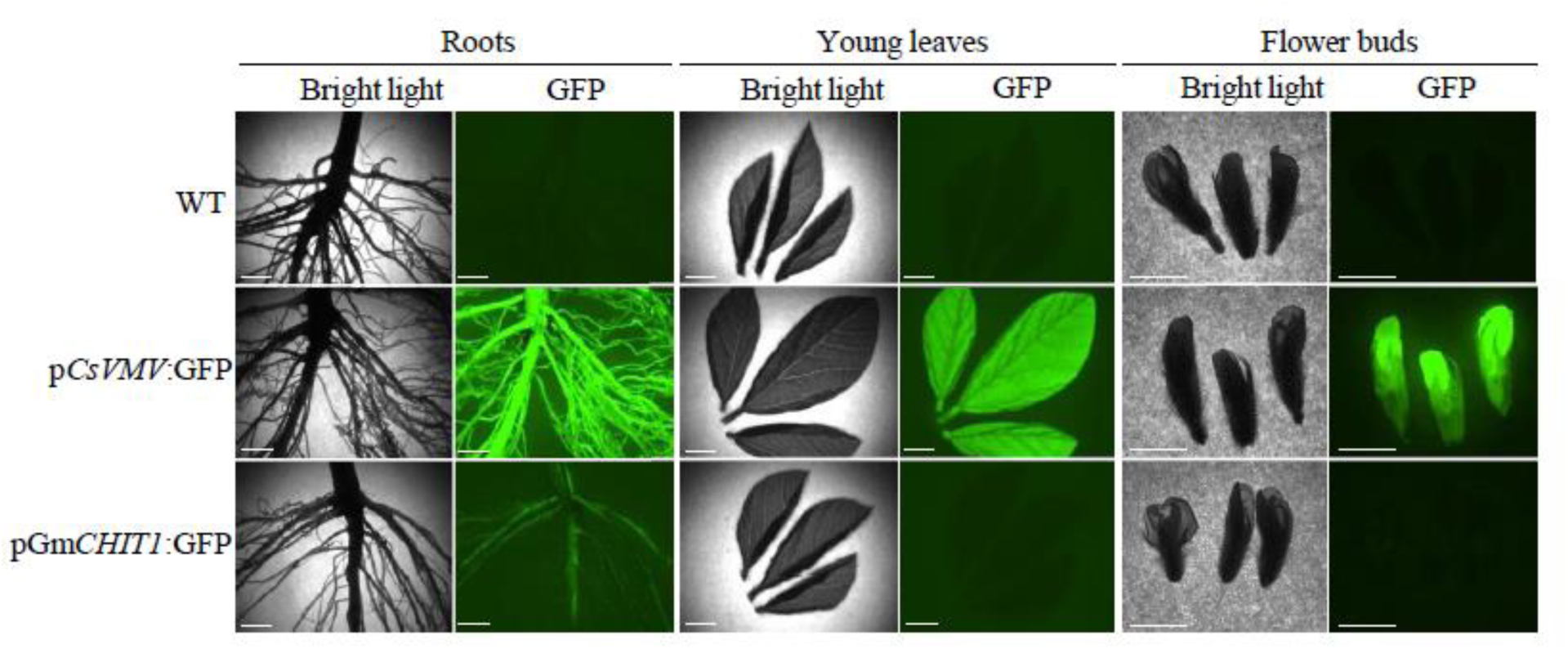
GFP activity mediated by *GmCHIT1* promoter (line 131 p*GmCHIT1*:GFP) in soybean tissues (roots, leaves, flower buds). Plants transformed with p*CsVMV*:GFP were used as a positive control. Bar-scales represent 5mm. Pictures were taken with a dissection scope (Leica Z16 APO) under GFP filter and bright light.

**Figure 5:**
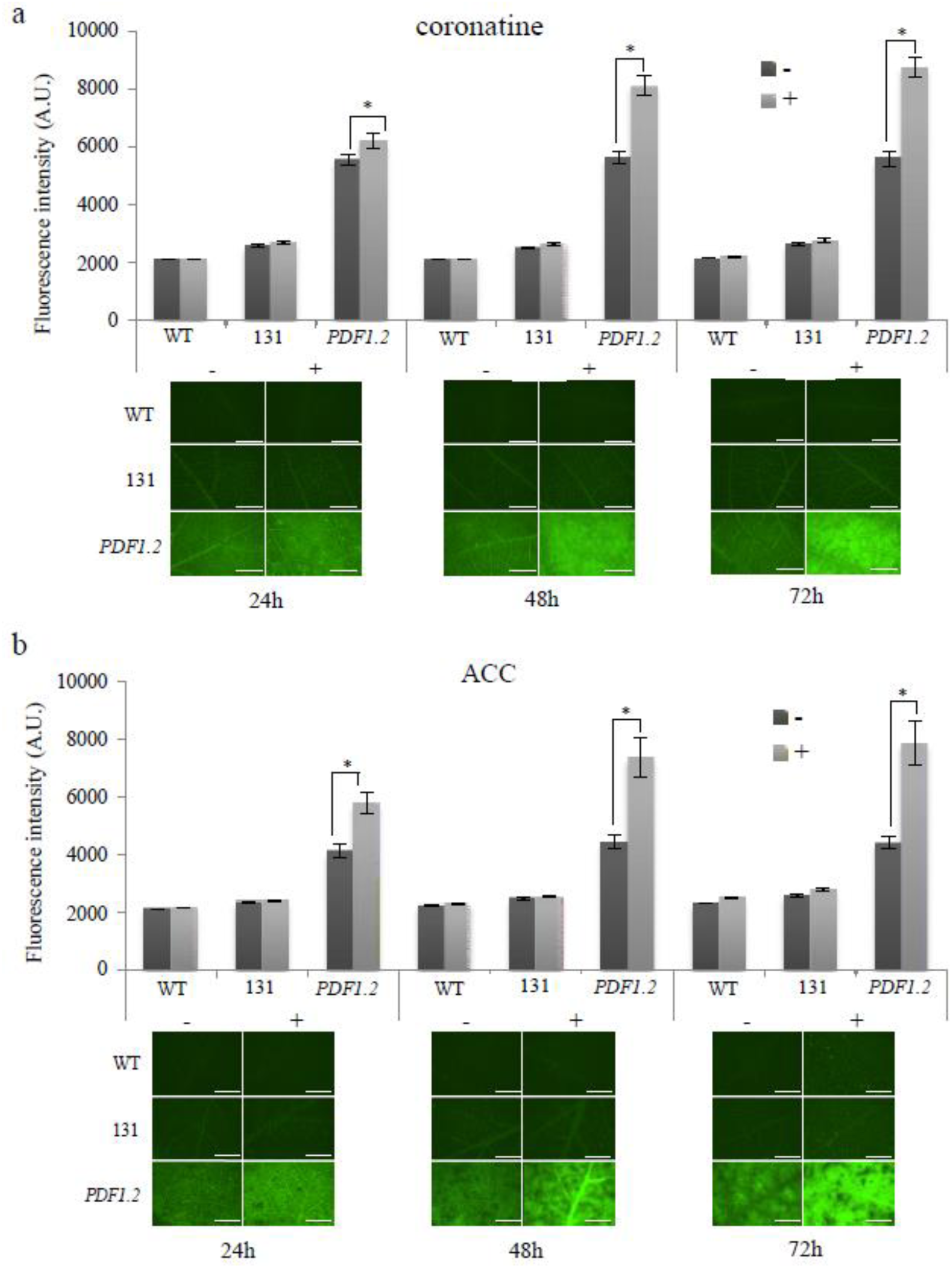
*GmCHIT1* promoter expression following hormonal treatments. GFP fluorescence observed in 131 line (p*GmCHIT1*:GFP), *PDF1.2* (p*PDF1.2*:GFP) or WT detached leaves following hormonal (+) or mock (-) treatments. Treatments were done with coronatine (a) and ACC (b). Graphics represent the mean ± standard errors of fluorescence intensity measured with MetaMorph software *via* grayscale value on 20 biological replicates. *: significant difference between treated (+) and untreated (-) leaves determined by a Student’s t-test (*p* < 0.05). Representative images of the observed fluorescence are shown under the graphs. Bar-scales represent 5mm. Observations were realized at 24, 48 and 72 hours after hormonal treatment with a dissection scope (Leica Z16 APO) under GFP filter.

**Figure 6:**
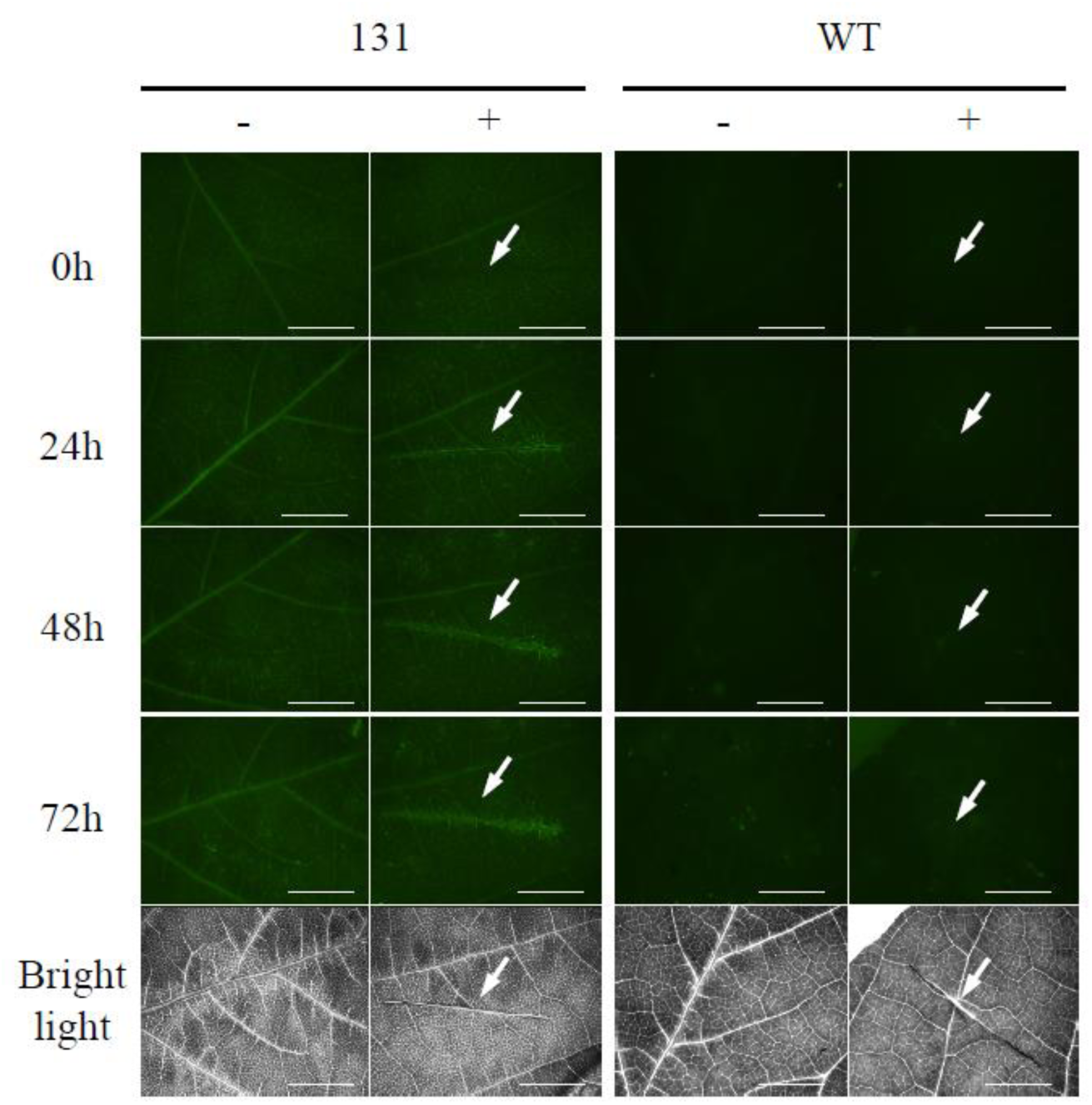
*GmCHIT1* promoter response after wounding. GFP fluorescence in wounded (+) or control (-) detached leaves from transgenic soybean (line 131 with the GFP fused to the *GmCHIT1* promoter) and WT plants. Bar-scale represents 5mm. Observations at 0, 24, 48 and 72 hours after wounding with a dissection-scope (Leica Z16 APO) under GFP filter and bright light. Arrows show the wounded part.

### Activity of the *GmCHIT1* promoter in response to hormone and wounding treatments

To evaluate the potential induction of the *GmCHIT1* promoter by other stimuli than fungal contamination, different hormonal treatments were performed on plants and the activity profile was evaluated in the line 131. For this, the plants were subjected to coronatine (methyl jasmonate analogue) and 1-aminocyclopropane-1-carboxylic acid (ACC, ethylene precursor) treatments. As *A. thaliana PDF1.2* promoter has been shown to be induced by jasmonate and ethylene ^22^, p*PDF1.2*:GFP soybean plants (named *PDF1.2*) were used as positive controls for these investigations. As expected, fluorescence strongly increased from 24 to 72 h after coronatine or ACC treatments (Figure 5a-b). In the case of the p*GmCHIT1*:GFP plants, fluorescence intensity did not change after coronatine or ACC spray (Figure 5a-b), suggesting that p*GmCHIT1* was not induced by these hormonal treatments. Fluorescence intensity remained also unchanged after salicylic acid (SA) exposure in p*GmCHIT1*:GFP plants (line 131) (additional file 8 Figure S7). As we had no functional control to evaluate the efficiency of this last treatment, the expression of three *PR* genes (*GmPR1, GmPR2* and *GmPR3*) ^38^, was followed by RT-qPCR in the leaves of plants from line 131. In our experimental conditions, only a low induction of *GmPR3* (2-fold change compared to mock) was detected in response to SA exposure (additional file 8 Figure S7). This last result did not allow to conclude on the efficiency of the treatment and consequently on the inducibility of p*GmCHIT1* by SA.

Lastly, *GmCHIT1* promoter response was monitored after mechanical wounding. A small GFP fluorescence was observed at 24 h post-wounding limited to the wounded area and still visible at 72 h after the injury (Figure 6). The *GmCHIT1* promoter appeared to be induced by wounding with no propagation to adjacent tissues.

## DISCUSSION

Today, biotechnology approaches can be considered to develop alternative strategies to control fungal diseases, and more specifically the rust pathogen *P. pachyrhizi.* In this context, many genes associated with disease resistance have been identified and proposed to develop transgenic plants capable of defending themselves against pathogens ^17,39,40^. To drive the expression of these genes only during pathogen infection, the use of pathogen-inducible promoters is recommended. Such promoters have been isolated in several plants from genes associated with defence response ^41^. This is the case for the barley germin-like GER4 promoter that controls the expression of a PR protein highly induced in response to biotrophic or necrotrophic pathogens ^20^. Nevertheless, the identification and characterization of such promoters in soybean is still limited ^24,42^. This work presents the identification of a soybean putative chitinase gene promoter (p*GmCHIT1)* and its activity profile in soybean plants. Several studies have shown that genes associated with defence response, such as PR genes, are found to be induced during soybean rust inoculation in both resistant and susceptible soybeans ^43^. Among them, the *GmCHIT1* gene coding for a putative chitinase was reported as upregulated during early (24 hpi) and later (10 dpi) stages of *P. pachyrhizi* infection. We investigated the expression profile of this gene during the infectious process of *P. pachyrhizi* on soybean plants and confirmed that the expression of this gene was detectable as early as 8 hpi, remained constant from 24 to 72 hpi and increased drastically at 10 dpi. Microscopic observations of the infectious development of *P. pachyrhizi* revealed that appressorium formation and rust penetration in plant tissues occur between six and twelve hours after urediniospore inoculation. Between 24 and 48 hpi, the fungus mainly forms haustoria and this differentiation step is rapidly followed by the fungal growth inside the host tissues^44^. Considering *GmCHIT1* expression, we can assume that it could be induced through a plant signal during the appressorium formation and/or fungal penetration, and its expression could be proportional to the quantity of mycelia developing inside the plant tissues.

Heterologous systems are often used to study gene expression, but results produced in these experiments are limited because promoter regulation may depend on the genetic background of the plant species under investigation ^45–47^. A transient system could allow a rapid investigation of a promoter’s activity, and the opportunity to select the smallest inducible promoter region. However, transient transformation of soybean is difficult to implement, and results are not still reproducible. We therefore generated stable transgenic soybean plants harbouring GFP placed under the control of the *GmCHIT1* promoter. This approach gave us the opportunity to highlight the local induction of the plant chitinase promoter in soybean cells surrounding fungal appressoria, the fungal penetration structures (Figure 3).

Mechanical injuries of plant tissues can provide an entrance for pathogen invasion. Therefore, several wound-induced genes are also involved in plant defence pathways against invading fungi ^48^. *P. pachyrhizi* penetrates directly the epidermal cells of the leaves rather than the stomata ^4^and this action leads to the collapse of the epidermal cells. In this particular case of interaction, it is not surprising to observe that p*GmCHIT1* is also induced after wounding. The pattern of p*GmCHIT1* response to wounding is similar to the one observed by Hernandez-Garcia and Finer in wounded soybean plants harbouring the transcriptional fusion of the GFP and *GmERF3* promoter ^42^. However, in the context of the Asian soybean rust infection, we cannot conclude that p*GmCHIT1* induction is the result of signalling associated solely with the tissue injury, the rust infection or both.

Some plant chitinase promoters have already been studied. Thus, the *BjChp* chitinase promoter of *Brassica juncea* has been reported to be induced by the pathogen *Alternaria brassicae*, jasmonic acid and wounding in *A. thaliana* ^49^. *BjChp* promoter activity was also observed surrounding the necrotic lesions at 48 hpi. Another chitinase promoter of *Phaseolus vulgaris* (*PvChi4*) promoter has been reported to be expressed in lateral roots and reproductive organs of non-stressed *A. thaliana* plants ^50^and it was also induced by heat treatment and UV light. Additionally, the promoter of the chitinase AtEP3, the closest *A. thaliana* orthologue of GmCHIT1, was shown to be early induced by *Xanthomonas campestris* at 1, 6 and 24 hpi but downregulated by wounding ^51,52^. These results highlight that chitinase promoters can be regulated by biotic or abiotic stresses or both.

Transcriptional regulation of defence genes under biotic stress is regulated by many *cis*-elements localized in the promoter ^21^. Among them, GCC-box and W-boxes have been shown to be inducible by pathogens and wounding ^21^. In the *ChiIV3* chitinase promoter of pepper, one W-box located in the - 712/-459 bp region was described as essential to trigger the induction after *Phytophthora capsici* contamination ^53^. W-box refers to the binding site of WRKY transcription factors ^33^, and in soybean, these regulators have been shown to be implicated in the response to *P. pachyrhizi* ^54^. In the *GmCHIT1* promoter, 21 W-boxes have been identified. In addition, 10 GT1-boxes and 5 MYB recognition elements have also been found. It has been demonstrated that GT1-boxes are responsible for the induction of defence genes by pathogen and high salinity stress, as it has been described for the soybean promoter of the calmodulin *SCaM-4*. ^55^. MYB recognition elements were found in defence gene promoters and could be implicated in response to abiotic stress and hormone treatment ^56^. Finally, two auxin and two gibberellic acid responsive elements have been found in the *GmCHIT1* promoter. These observations suggest that p*GmCHIT1* could be potentially activated by these hormones. It would be interesting to investigate whether the *cis*-regulating elements found in the *GmCHIT1* promoter are essential and sufficient to trigger a response to *P. pachyrhizi*.

Fungal infection can induce different plant hormone pathways depending on the lifestyle of the pathogen. It is well-admitted that salicylate signalling is implicated in defence against biotrophic fungi and jasmonate together with ethylene participate in the defence against necrotrophic fungi ^57^. However, a study of non-host interaction between *P. pachyrhizi* and *A. thaliana* has revealed that despite the biotrophic lifestyle of *P. pachyrhizi*, the pathogen activates marker genes of necrotrophic infection ^58^. It has been suggested that the fungus would mimic a necrotrophic behaviour at the initial stage of infection to promote its development inside the host tissues ^59^. In this context, one would expect *P. pachyrhizi* development to induce the jasmonic acid or ethylene pathway at early time-points after inoculation and salicylic acid-related genes at later times. However, expression data during the early and late stages of *P. pachyrhizi* development in soybean did not reveal clear evidence of activation of either the salicylate, jasmonic acid or ethylene pathway ^30,43,59^. Nevertheless, it was surprising to observe that the *GmCHIT1* promoter was not induced by any hormonal treatments assessed in our study. Indeed, several PR proteins have been shown to be activated by plant hormones ^60^. For instance, a chitinase from rice has been reported to be induced by jasmonic acid and ethylene 48 h post-treatment ^61^. However, unlike in Mazarei *et al.* ^62^, in our experimental conditions, *GmPR1* was not induced after salicylic acid treatment and only a slight induction of *GmPR3* was observed. It is unclear at this stage whether the results reflect a lack of efficacy of salicylic acid treatment or an insensitivity of p*GmCHIT1* to this hormone.

Basal *GmCHIT1* promoter activity in non-contaminated soybean tissues was also investigated. Visualization of GFP expression revealed that p*GmCHIT1* was expressed in the veins of fully developed leaves and in roots but not in young leaves and flowers. Roots are permanently exposed to soil pathogens that can penetrate the tissues because of micro-wounds and the absence of lignified barriers ^63^. This basal expression level in different soybean tissues/organs together with the induction under rust attack might reflect the potential roles of this chitinase in physiological processes of growth and development as much as in pathogen protection. Nevertheless, despite the basal expression of this promoter, it can be considered as an interesting tool to monitor expression of defence genes. Indeed, in addition to its inducible characteristic, we observed that its basal expression in soybean tissues remained lower than the constitutive expression of CsVMV promoter. This makes p*GmCHIT1* a prime candidate compared to constitutive promoters.

## CONCLUSIONS

Promoters are the primary regulators of gene expression at the transcriptional level and are considered as key elements to control genes of interest in transgenic organisms. Pathogen inducible promoters allowing transgene expression only when and where it is needed, are interesting tools for the development of biotechnological approaches to control bacterial or fungal diseases. In this study, we identified p*GmCHIT1*, a promoter of a soybean predicted chitinase gene expressed during the first hours of the Asian soybean rust disease. Moreover, this promoter is reported as locally activated by *P. pachyrhizi* on plant tissue. To our knowledge, p*GmCHIT1* is the only promoter isolated to date in soybean with such traits. These characteristics suggest that it could be therefore considered as a candidate for driving defence genes in genetically engineered soybean.

## METHODS

### Construction of the transformation vectors

The GFP reporter gene ^64^ was amplified by PCR with primers gfp-F/gfp-R (see additional file 9: Table S2) and cloned downstream of the CsVMV promoter from Cassava Vein Mosaic Virus ^65^. The *PDF1.2* promoter from *Arabidopsis thaliana* ^22^ was amplified by PCR using primers pdf1.2-F/ pdf1.2-R (see additional file 9: Table S2) and cloned to drive the expression of the GFP-encoding sequence. Upstream of the *Glyma.11G124500* gene-encoding sequence (based on *G. max* genome sequence from https://phytozome.jgi.doe.gov/pz/portal.html), a 3454 bp segment considered as part of the *GmCHIT1* promoter was synthesized by Eurofins genomic (Germany). The promoter was then cloned to drive the expression of the GFP-encoding gene. Each GFP construct was transferred to *A. tumefaciens* strain LBA4404. In all vectors, the HPPD (hydroxyphenylpyruvate dioxygenase) gene driven by the 35S promoter was used as a selectable marker for soybean transformation ^66^.

### Soybean cultivation

Seeds of soybean cultivar Thorne, susceptible to *P. pachyrhizi*, were sown in pots containing SteckMedium substrate (Klasmann-Deilmann GmbH, Germany) for germination. After 3 weeks, the plants were transferred into larger pots for development and eventually seed production. Greenhouse conditions were as follows: temperature of 24 °C day/22 °C night with a photoperiod of 16 h of day under a light intensity of 270 μE.m^-2^.s^-1^ and 70% relative humidity.

### Soybean transformation

Seeds were surface sterilized for 24 h in a desiccator by chlorine gas generated with a mixture of 150 ml Domestos containing 4.5% NaClO w/w (Unilever) and 5 ml of HCl (37%). Sterile seeds were then hydrated overnight in sterile deionized water. Cotyledons of germinated seeds were dissected by removing the seed coat and by splitting the seeds into 2 halves using a scalpel blade. The half-seeds were immersed for 30 min in 10% W/V Gamborg’s medium (Gamborg *et al*., 1968) containing 30 g/l sucrose, 7.4 μM BAP (6-benzylaminopurine), 0.7 μM GA3 (gibberellic acid A3), 3.3 mM cysteine, 1 mM dithiothreitol, 200 μM acetosyringone, 20 mM MES, pH 5.4 and the bacterium *Agrobacterium tumefaciens* at a final OD_600nm_ of 0.8. Next, cotyledons were transferred to Petri dishes, adaxial side down, onto 3 layers of Whatman ® paper pre-soaked with 10 ml of Gamborg’s medium. Plates were transferred to a tissue culture room for 5 days at 24°C, 16 h light (180 μE.m^-2^.s^-1^) and 75% relative humidity. Shoots were induced by transferring the cotyledons to full-strength Gamborg’s medium containing 30 g/l sucrose, 7.4 μM BAP, 3 mM MES pH 5.6 and 8 g/l noble agar. Antibiotics ticarcillin (50 mg/l), cefotaxime (50 mg/l), vancomycin (50 mg/l) and the herbicide Tembotrione™ (0.2 mg/l) used as selectable marker were added after autoclaving. After one month on the shoot induction medium, white shoots were removed and cotyledons were transferred on a shoot elongation medium containing Murashige & Skoog (MS) salts ^67^, 3.2 g/l Gamborg’s vitamins, 30 g/l sucrose, 100 mg/l pyroglutamic acid, 50 mg/l asparagine, 0.28 μM zeatin riboside, 0.57 μM indol-3-acetic acid, 14.8 μM GA3, 3 mM MES, pH 5.6 and 8 g/l noble agar. Antibiotics and the herbicide were kept at the same concentrations previously described. After one month, elongated shoots were cut and transferred to a rooting medium consisting of half-strength MS salts, half-strength B5 vitamins, 15 g/l sucrose, and 8 g/l noble agar. The same antibiotics as previously described were added after autoclaving, but the selectable marker was omitted. When roots were sufficiently developed, the shoots were individually transplanted to a greenhouse and cultivated using the conditions previously described.

### Characterization of transgenic plants

Regenerated T0 events were confirmed for the presence of the selectable marker gene with an HPPD lateral flow test (AMAR Immunodiagnostics) using the experimental instructions recommended by the provider. To pick up T1 HPPD/GFP-positive events, germinated seeds were watered with an 8‰ solution of the herbicide Isoxaflutole™ to eliminate null segregant plants. Plants showing no herbicide symptoms were subsequently tested for GFP fluorescence and used for further analysis. Homozygous single-locus plants were selected either in T1 or T2 segregating generations by ddPCR analysis. T1 or T2 plants were used depending on the availability of the material.

### Fungal contamination of soybean plants

A dehydrated stock of spores of *P. pachyrhizi* stored in liquid nitrogen (isolate MG2006, Mato Grosso, Brazil 2006) was used as a routine source of inoculum. Twenty-four hours before plant inoculation, cryo-tubes were opened and placed in a controlled growth chamber (20°C, dark, 70% relative humidity) to slowly rehydrate the spores. The spores were finally suspended in sterilized water containing 0.01% Tween 20 to reach a final concentration of 100,000 spores/ml. Three-week-old soybean plants were sprayed with the spores until run-off and incubated in a growth chamber (temperature 24°C, dark, 100% relative humidity) for 24 h before being transferred to a developing chamber (temperature of 24°C, 16 h light/8 h night, light intensity 15 μE.m^-2^.s^-1^ and 80% relative humidity). All experiments were conducted according to the recommendations of the French biosafety agency (Haut Conseil des Biotechnologies).

### Treatment of detached soybean leaves

First and second trifoliate leaves of 6-week-old plants were excised and transferred to layers of Whatman® paper wetted with 6 ml of sterile distilled water. Leaf petioles were wrapped with water-soaked cotton to increase organ survival. Different hormone treatments were conducted by spraying leaves with either 20 mM of ACC (ethylene precursor) or 2.5 mM solution of salicylic acid (SA) in sterile water or 0.25 mM of coronatine (methyl jasmonate analogue) in 1% EC premix solution (phenyl sulfonate 5%, emulsogen EL360 7%, isophorone 40% and methyloleate 48%). Sterile distilled water was used as mock for ACC and SA treatments, and 1% EC premix was used as mock for coronatine spray. Leaf wounding was realized with a sterile scalpel blade. After the different treatments, the leaves were incubated in the same growth chamber used for soybean transformation. Macroscopic observations and fluorescence intensity measurement were performed at 24, 48 and 72 h post-treatment.

### Expression profiling by quantitative PCR analysis

Samples were composed of four foliar discs from leaves of a soybean plant, and three independent biological replicates were performed. Total RNA was extracted using the RNeasy^®^ Plant Mini Kit (Qiagen, Netherlands) and purified with the TURBO DNA-*free*™ Kit (Invitrogen, Carlsbad, CA). DNA-free total RNA (1 µg) was used to synthetize cDNA with the ThermoScript™ RT-PCR System kit (Invitrogen, Carlsbad, CA) according to the manufacturer’s recommendations. For RT-qPCR, 0.02 µg of cDNA was used in a 20 µl reaction containing 10 µl of SsoAdvanced™ Universal SYBR^®^ Green Supermix (Bio-Rad, US), 6 µM of forward and reverse primers and 3 µl of RNAse-free water. RT-qPCR was performed using the LightCycler® 480. The thermocycling conditions were followed as recommended by the supplier. The expression of the chitinase gene was determined after soybean rust inoculation by using specific primers (see additional file 9: Table S2). The genes coding for actin (GenBank: NM_001289231.2) and a hypothetical protein (GenBank: BE330043) ^65^ (primer sequences in additional file 9: Table S2) were used as endogenous reference genes for normalization ^68^ using the Ct value method. Specific primers of *GmPR1* (GenBank: BU5773813), *GmPR2* (GenBank: M37753) and *GmPR3* (GenBank: AF202731) were used to determine the expression of those *PR* genes after salicylic acid treatment. In this case, the genes coding for actin and an elongation factor (GenBank: NM_001249608.2) were used for normalization (see additional file 9: Table S2) with the Ct value method.

### Western blot analysis

Leaf samples from wild-type (WT) plants and plants from line 131 were harvested 72 h after *P. pachyrhizi* contamination or mock treatment. Proteins were extracted from four foliar discs of the same soybean plant with 250 µl of extraction buffer (Tris-Hcl 100 mM, NaCl 100 mM, DTT 0.04%) and placed on ice for 10 min before centrifugation at 4°C for 10 min. The protein concentration was determined with the Bradford method using the Bio-Rad Protein assay dye reagent solution. For denaturation, 1 volume of Laemmli buffer (Bio-Rad, US) was added to 1 volume of extracted proteins (30 µg). The mixture was kept for 5 min at 95°C and 5 min on ice before loading on a TGX 4-20% Strainfree (Bio-Rad US) gel immersed in TGS 1X buffer. After migration, separated proteins were transferred onto a membrane by using Trans-Blot® Turbo™ Midi Nitrocellulose Transfer Packs (Bio-Rad) and the TransBlot Turbo device (Bio-Rad, US). Membrane blocking and incubation with the antibodies were performed as suggested by the provider. GFP antibodies (Sigma) and Immun-Star Goat Anti-Rabbit (GAR)-HRP Conjugate antibody were used. Antibody detection was realized with the Clarity™ Western ECL (Bio-Rad, US) kit following the supplier’s instructions. Finally, the ChemiDoc™ Touch camera (Bio-Rad, US) was used to record the results.

### Visualization of GFP expression

GFP fluorescence was analysed with a Leica Z16 APO A dissection scope equipped with a GFP filter. For the detection of fluorescence after rust inoculation, the parameters were set as follows: camera lens 1 x, magnification 115 x, gain 2 and exposure time 500 ms. For detection of the GFP fluorescence in the different soybean tissues without infection, the camera lens was set at camera lens 0.5 x, magnification at 6.95 x for roots and young trifoliate leaves and 15 x for flowers, gain 3, exposure time 500 ms. For hormonal treatments and wounding, the following parameters were used: camera lens 1 x, magnification 6.95 x, gain 3 and exposure time 1 s. Fluorescence intensity measurement was performed using MetaMorph software *via* greyscale value.

### Confocal microscopy

Leaf samples of soybean line 131 expressing the transcriptomic fusion p*GmCHIT1*:GFP were harvested 24 h post-inoculation. The samples were first stained in an aqueous calcofluor white solution (0.01 mg/ml) for 5 min before being washed 3 times in water for 5 min. Samples were mounted in water under slides (VWR® microscope slides: ground edges 45°, 76 x 26 mm) and cover glass (VWR® cover glass: 22 x 32 mm). Observations were conducted with a ZEISS LSM 800 microscope using the 10x objective. To visualize GFP fluorescence, a 487 nm wavelength laser was used for excitation and light emission was captured at 560 nm. For the imaging of calcofluor fluorescence, light excitation was set at a wavelength of 400 nm and emission was captured at 487 nm.

## ETHICS APPROVAL AND CONSENT TO PARTICIPATE

Not applicable

## CONSENT FOR PUBLICATION

Not applicable

## AVAILABILITY OF DATA AND MATERIAL

All the data and material generated are BASF property.

## AUTHOR’S CONTRIBUTIONS

NP, BP, SD and LC conceived and designed the experiments. SP performed the gene expression analyses, LG the soybean transformations and LC the rest of the experiments. NP, BP, SD, CS, SP and LC analysed the data. NP, BP, SD and LC wrote the paper. All the authors have read and approved the manuscript.

## ACKNOWLEDGEMENTS

We gratefully acknowledge Pr. Ulrich Schaffrath, Dr. Marc-Henri Lebrun, Dr. Frank Meulewaeter and Dr. Florent Villiers for their advices and discussions as well as for their comments on the manuscript. We also thank Didier Joiris for the soybean culture and its excellent work in greenhouse and Laura Velazquez for her contribution on the project.

