## supplementary data for "Identification and characterization of a new soybean promoter induced by *Phakopsora pachyrhizi*, the causal agent of Asian soybean rust"

### 1 ADDITIONAL FILES

2

| Gene | Gene_Description | Function_2010 | Rank 2010 | Rank 2011 |
| --- | --- | --- | --- | --- |
| Glyma.02G145200 | E3 ubiquitin ligase involved in syntaxin degradation | Cell Growth & Division | 213 | 1297 |
| Glyma.12G022500 | E3 ubiquitin ligase involved in syntaxin degradation | Cell Growth & Division | 199 | 542 |
| Glyma.10G275200 | Uncharacterized conserved protein | Cell Growth & Division | 260 | 989 |
| Glyma.10G212900 | Translation elongation factor EF-1 alpha/Tu | Cell Growth & Division | 141 | 53 |
| Glyma.08G358300 | 26S proteasome regulatory complex, subunit PSMD10 | Cell Structure | 467 | 1930 |
| Glyma.12G053900 | Beta-glucosidase, lactase phlorizinhydrolase, and related proteins | Cell Structure | 259 | 837 |
| Glyma.14G217100 | Histones H3 and H4 | Cell Structure | 492 | 3113 |
| Glyma.18G288600 | Phospholipase D1 | Cell Structure | 490 | 906 |
| Glyma.02G093500 | Phospholipase D1 | Cell Structure | 258 | 2633 |
| Glyma.17G008000 | Predicted membrane protein, contains DoH and Cytochrome b-561/ferric reductase transmembrane domains | Cell Structure | 22 | 186 |
| Glyma.07G266300 | Predicted membrane protein, contains DoH and Cytochrome b-561/ferric reductase transmembrane domains | Cell Structure | 294 | 1663 |
| Glyma.13G091000 | Tetraspanin family integral membrane protein | Cell Structure | 164 | 1693 |
| Glyma.17G159300 | Uncharacterized membrane protein | Cell Structure | 562 | 4352 |
| Glyma.14G004300 | Predicted NUDIX hydrolase FGF-2 and related proteins | Cell Structure | 15 | 61 |
| Glyma.02G126800 | Protein transporter of the TRAM (translocating chain-associating membrane) superfamily | Cell Structure | 401 | 189 |
| Glyma.02G107500 | Membrane associated zinc finger | Cell Structure | 313 | 4512 |
| Glyma.19G054700 | Leucine rich repeat protein | Disease & Defence | 266 | 134 |
| Glyma.05G082200 | Leucine rich repeat protein | Disease & Defence | 379 | 466 |
| Glyma.17G179200 | Leucine rich repeat protein | Disease & Defence | 595 | 439 |
| Glyma.05G055000 | Copper chaperone for superoxide dismutase | Disease & Defence | 198 | 2136 |
| Glyma.15G062400 | Defence-related protein containing SCP domain | Disease & Defence | 14 | 1 |
| Glyma.15G062500 | Defence-related protein containing SCP domain | Disease & Defence | 11 | 22 |
| Glyma.10G152200 | Ferric reductase, NADH/NADPH oxidase and related proteins | Disease & Defence | 288 | 1302 |
| Glyma.19G233900 | Ferric reductase, NADH/NADPH oxidase and related proteins | Disease & Defence | 479 | 2405 |
| Glyma.04G254100 | Galactosyltransferases | Disease & Defence | 422 | 1897 |
| Glyma.13G346700 | Predicted chitinase | Disease & Defence | 2 | 5 |
| Glyma.11G124500 | Predicted chitinase | Disease & Defence | 4 | 40 |
| Glyma.20G150200 | dioxygenase | Metabolism | 545 | 1002 |
| Glyma.10G244100 | dioxygenase | Metabolism | 542 | 1568 |
| Glyma.04G232200 | AAA+type ATPase | Energy | 183 | 3779 |
| Glyma.19G018600 | AAA+type ATPase | Energy | 145 | 157 |
| Glyma.07G052400 | Cytochrome b5 | Energy | 454 | 1188 |
| Glyma.16G019900 | D-arabinono-1,4-lactone oxidase | Energy | 466 | 34 |
| Glyma.06G107200 | Glutaredoxin and related proteins | Energy | 189 | 493 |
| Glyma.01G210400 | Lactate dehydrogenase | Energy | 19 | 556 |
| Glyma.05G058100 | Lactate dehydrogenase | Energy | 443 | 3469 |
| Glyma.17G128000 | Malate synthase | Energy | 345 | 52 |
| Glyma.11G179300 | NADH-dehydrogenase (ubiquinone) | Energy | 90 | 8 |
| Glyma.06G266700 | Thioredoxin | Energy | 123 | 2951 |
| Glyma.08G285900 | Thioredoxin, nucleoredoxin and related proteins | Energy | 300 | 362 |
| Glyma.04G021000 | Thioredoxin, nucleoredoxin and related proteins | Energy | 565 | 1096 |
| Glyma.08G277000 | Transketolase | Energy | 84 | 214 |
| Glyma.08G162200 | Uncharacterized high-glucose-regulated protein | Energy | 548 | 3069 |
| Glyma.03G246700 | Voltage-gated shaker-like K channel, subunit beta/KCNAB | Energy | 575 | 1710 |
| Glyma.17G076600 | SNAP-25 (synaptosome-associated protein) component of SNARE complex | Intracellular Traffic | 201 | 754 |
| Glyma.05G023100 | SNAP-25 (synaptosome-associated protein) component of SNARE complex | Intracellular Traffic | 257 | 2916 |
| Glyma.13G180100 | AAA+type ATPase | Intracellular Traffic | 444 | 3780 |
| Glyma.04G060700 | Endosomal membrane proteins, EMP70 | Intracellular Traffic | 588 | 1952 |
| Glyma.05G047800 | Exocyst component protein and related proteins | Intracellular Traffic | 468 | 1163 |
| Glyma.03G174300 | Exocyst component protein and related proteins | Intracellular Traffic | 321 | 280 |
| Glyma.15G013900 | G-protein beta subunit | Intracellular Traffic | 61 | 621 |
| Glyma.17G146100 | GTPase Rab1/YPT1, small G protein superfamily, and related GTP-binding proteins | Intracellular Traffic | 473 | 5641 |
| Glyma.08G239900 | Phosphatidylinositol transfer protein SEC14 and related proteins | Intracellular Traffic | 346 | 3260 |
| Glyma.18G013100 | Predicted E3 ubiquitin ligase | Intracellular Traffic | 517 | 743 |
| Glyma.19G051500 | Predicted E3 ubiquitin ligase | Intracellular Traffic | 442 | 5589 |
| Glyma.15G069100 | Secretory carrier membrane protein | Intracellular Traffic | 104 | 5353 |
| Glyma.02G195400 | SNARE protein Syntaxin 1 and related proteins | Intracellular Traffic | 166 | 755 |
| Glyma.02G195300 | SNARE protein Syntaxin 1 and related proteins | Intracellular Traffic | 102 | 151 |
| Glyma.13G136600 | SNARE protein YKT6, synaptobrevin/VAMP superfamily | Intracellular Traffic | 581 | 5188 |
| Glyma.08G254500 | 6-phosphogluconate dehydrogenase | Metabolism | 320 | 812 |
| Glyma.10G200700 | Acetylglucosaminyltransferase EXT2/exostosin 2 | Metabolism | 428 | 576 |
| Glyma.14G042000 | Alcohol dehydrogenase, class III | Metabolism | 458 | 1048 |
| Glyma.02G034000 | Aldehyde dehydrogenase | Metabolism | 82 | 727 |
| Glyma.05G231800 | Aldehyde dehydrogenase | Metabolism | 350 | 1225 |
| Glyma.18G285800 | Aldo/keto reductase family proteins | Metabolism | 417 | 816 |
| Glyma.02G307300 | Aldo/keto reductase family proteins | Metabolism | 34 | 2205 |
| Glyma.03G148300 | Arylacetamide deacetylase | Metabolism | 299 | 1555 |
| Glyma.01G239600 | Arylacetamide deacetylase | Metabolism | 48 | 5047 |
| Glyma.18G061100 | Asparagine synthase (glutamine-hydrolysing) | Metabolism | 36 | 60 |
| Glyma.12G005100 | Beta-fructofuranosidase (invertase) | Metabolism | 160 | 1255 |
| Glyma.08G028200 | Choline phosphate<br>cytidyltransferase/Predicted CDP-ethanolamine synthase | Metabolism | 606 | 5615 |
| Glyma.11G091400 | Copper chaperone | Metabolism | 148 | 2 |
| Glyma.10G172700 | Cystathionine beta-lyases/cystathionine<br>gamma-synthases | Metabolism | 88 | 204 |
| Glyma.08G140600 | Cytochrome P450 CYP2 subfamily | Metabolism | 163 | 33 |
| Glyma.13G285300 | Cytochrome P450 CYP2 subfamily | Metabolism | 165 | 143 |
| Glyma.11G062500 | Cytochrome P450 CYP2 subfamily | Metabolism | 85 | 254 |
| Glyma.13G173500 | Cytochrome P450 CYP2 subfamily | Metabolism | 170 | 287 |
| Glyma.07G089800 | Cytochrome P450 CYP2 subfamily | Metabolism | 175 | 376 |
| Glyma.15G156100 | Cytochrome P450 CYP2 subfamily | Metabolism | 63 | 378 |
| Glyma.13G068800 | Cytochrome P450 CYP2 subfamily | Metabolism | 211 | 608 |
| Glyma.03G143700 | Cytochrome P450 CYP2 subfamily | Metabolism | 597 | 1282 |
| Glyma.07G202300 | Cytochrome P450 CYP2 subfamily | Metabolism | 274 | 1570 |
| Glyma.18G080400 | Cytochrome P450 CYP2 subfamily | Metabolism | 452 | 2347 |
| Glyma.03G031000 | Cytochrome P450 CYP2 subfamily | Metabolism | 614 | 3714 |
| Glyma.13G262000 | Cytochrome P450 CYP4/CYP19/CYP26<br>subfamilies | Metabolism | 327 | 455 |
| Glyma.04G035000 | Cytochrome P450 CYP4/CYP19/CYP26<br>subfamilies | Metabolism | 485 | 2352 |
| Glyma.11G215700 | Diadenosine and diphosphoinositol<br>polyphosphate phosphohydrolase | Metabolism | 240 | 1951 |
| Glyma.19G050100 | GDP-mannose pyrophosphorylase | Metabolism | 456 | 1308 |
| Glyma.16G063200 | Glucose-6-phosphate 1-dehydrogenase | Metabolism | 393 | 1600 |
| Glyma.11G213000 | Glutamate decarboxylase/sphingosine<br>phosphate lyase | Metabolism | 75 | 344 |
| Glyma.13G233300 | Glutamate-gated kainate-type ion channel<br>receptor subunit GluR5 and related<br>subunits | Metabolism | 267 | 615 |
| Glyma.11G215500 | Glutamine synthetase | Metabolism | 157 | 152 |
| Glyma.18G041100 | Glutamine synthetase | Metabolism | 101 | 436 |
| Glyma.12G207400 | Glutaredoxin and related proteins | Metabolism | 78 | 1601 |
| Glyma.10G139400 | Haloacid dehalogenase-like hydrolase | Metabolism | 448 | 3704 |
| Glyma.07G048900 | Hydroxyindole-O-methyltransferase and<br>related SAM-dependent methyltransferases | Metabolism | 195 | 10 |
| Glyma.09G281900 | Hydroxyindole-O-methyltransferase and<br>related SAM-dependent methyltransferases | Metabolism | 619 | 67 |
| Glyma.01G225200 | Long-chain acyl-CoA synthetases (AMP-forming) | Metabolism | 488 | 717 |
| Glyma.17G044300 | Lysine-ketoglutarate<br>reductase/saccharopine dehydrogenase | Metabolism | 569 | 182 |
| Glyma.10G291400 | Lysophospholipase | Metabolism | 344 | 583 |
| Glyma.13G354900 | NADP+dependent malic enzyme | Metabolism | 463 | 3131 |
| Glyma.19G011700 | Peroxidase/oxygenase | Metabolism | 27 | 167 |
| Glyma.02G309900 | Phosphorylcholine<br>transferase/cholinephosphate<br>cytidyltransferase | Metabolism | 533 | 1607 |
| Glyma.13G210000 | Predicted hydrolase related to diene lactone hydrolase | Metabolism | 121 | 20 |
| Glyma.19G099400 | Predicted hydrolase/acyltransferase (alpha/beta hydrolase superfamily) | Metabolism | 285 | 2048 |
| Glyma.01G205900 | Predicted lipase | Metabolism | 395 | 4099 |
| Glyma.02G148400 | Predicted PhzC/PhzF-type epimerase | Metabolism | 210 | 97 |
| Glyma.16G042000 | Reductases with broad range of substrate specificities | Metabolism | 203 | 446 |
| Glyma.17G067800 | Trehalose-6-phosphate synthase component TPS1 and related subunits | Metabolism | 108 | 284 |
| Glyma.09G030300 | UDP-glucose 4-epimerase/UDP-sulfoquinovose synthase | Metabolism | 370 | 3596 |
| Glyma.08G244800 | UDP-glucuronosyl and UDP-glucosyl transferase | Metabolism | 380 | 36 |
| Glyma.08G244700 | UDP-glucuronosyl and UDP-glucosyl transferase | Metabolism | 357 | 64 |
| Glyma.02G104300 | UDP-glucuronosyl and UDP-glucosyl transferase | Metabolism | 505 | 137 |
| Glyma.11G000500 | UDP-glucuronosyl and UDP-glucosyl transferase | Metabolism | 335 | 768 |
| Glyma.19G243300 | Uncharacterized conserved protein | Metabolism | 559 | 3696 |
| Glyma.06G135100 | Predicted RNA-binding protein SEB4 (RRM superfamily) | Post-Transcription | 514 | 400 |
| Glyma.15G154000 | Cullins | Protein Destination & Storage | 451 | 2137 |
| Glyma.04G049900 | Asparaginyl peptidases | Protein Destination & Storage | 593 | 1011 |
| Glyma.12G179800 | Aspartyl protease | Protein Destination & Storage | 177 | 831 |
| Glyma.01G163000 | Aspartyl protease | Protein Destination & Storage | 152 | 1018 |
| Glyma.05G064700 | Bax-mediated apoptosis inhibitor TEGT/BI-1 | Protein Destination & Storage | 475 | 2019 |
| Glyma.15G177800 | Cysteine proteinase Cathepsin L | Protein Destination & Storage | 360 | 107 |
| Glyma.07G140000 | Glutathione S-transferase | Protein Destination & Storage | 205 | 163 |
| Glyma.16G130700 | Hydrolytic enzymes of the alpha/beta hydrolase fold | Protein Destination & Storage | 394 | 1588 |
| Glyma.06G120600 | Leucine rich repeat proteins, some proteins contain F-box | Protein Destination & Storage | 520 | 352 |
| Glyma.17G049600 | Leucine rich repeat proteins, some proteins contain F-box | Protein Destination & Storage | 605 | 884 |
| Glyma.06G116300 | Mitochondrial inner membrane protease, subunit IMPI | Protein Destination & Storage | 303 | 1967 |
| Glyma.11G018500 | Mitochondrial solute carrier protein | Protein Destination & Storage | 430 | 3999 |
| Glyma.13G289600 | Molecular chaperone (DnaJ superfamily) | Protein Destination & Storage | 287 | 293 |
| Glyma.04G190800 | Peptide methionine sulfoxide reductase | Protein Destination & Storage | 132 | 71 |
| Glyma.06G317800 | Predicted membrane protein | Protein Destination & Storage | 498 | 24 |
| Glyma.15G156200 | Serine carboxypeptidases (lysosomal cathepsin A) | Protein Destination & Storage | 265 | 26 |
| Glyma.13G117700 | Ubiquitin and ubiquitin-like proteins | Protein Destination & Storage | 129 | 285 |
| Glyma.13G117900 | Ubiquitin and ubiquitin-like proteins | Protein Destination & Storage | 531 | 2159 |
| Glyma.15G091400 | Ubiquitin-like protein | Protein Destination & Storage | 566 | 724 |
| Glyma.04G238800 | Ubiquitin-protein ligase | Protein Destination & Storage | 438 | 5484 |
| Glyma.07G060900 | Translation initiation factor 1A (eIF-1A) | Protein Synthesis | 564 | 1616 |
| Glyma.02G236500 | Cytochrome P450 CYP2 subfamily | Secondary Metabolism | 497 | 5406 |
| Glyma.05G147000 | O-methyltransferase | Secondary Metabolism | 46 | 9 |
| Glyma.02G239500 | Acyl-CoA synthetase | Secondary Metabolism | 91 | 115 |
| Glyma.06G202300 | Cytochrome P450 CYP2 subfamily | Secondary Metabolism | 301 | 3738 |
| Glyma.09G269500 | Flavonol reductase/cinnamoyl-CoA reductase | Secondary Metabolism | 107 | 161 |
| Glyma.09G269600 | Flavonol reductase/cinnamoyl-CoA reductase | Secondary Metabolism | 83 | 51 |
| Glyma.02G158700 | Flavonol reductase/cinnamoyl-CoA reductase | Secondary Metabolism | 383 | 90 |
| Glyma.20G213700 | Hydroxyindole-O-methyltransferase and related SAM-dependent methyltransferases | Secondary Metabolism | 196 | 87 |
| Glyma.07G168500 | Iron/ascorbate family oxidoreductases | Secondary Metabolism | 232 | 196 |
| Glyma.15G223900 | NADH:flavin oxidoreductase/12-oxophytodienoate reductase | Secondary Metabolism | 122 | 314 |
| Glyma.03G181600 | Phenylalanine and histidine ammonia-lyase | Secondary Metabolism | 126 | 2623 |
| Glyma.12G059100 | Reductases with broad range of substrate specificities | Secondary Metabolism | 47 | 218 |
| Glyma.14G004500 | Predicted unusual protein kinase | Signal Transduction | 599 | 923 |
| Glyma.05G082400 | Apoptotic ATPase | Signal Transduction | 280 | 75 |
| Glyma.03G138000 | Ca<sup>2+</sup> dependent protein kinase, EF-Hand protein superfamily | Signal Transduction | 297 | 4622 |
| Glyma.19G257800 | Ca<sup>2+</sup> binding protein (centrin/caltractin), EF-Hand superfamily protein | Signal Transduction | 339 | 437 |
| Glyma.14G081300 | Ca<sup>2+</sup> independent phospholipase A2 | signal Transduction | 66 | 117 |
| Glyma.04G245000 | Calmodulin and related proteins (EF-Hand superfamily) | Signal Transduction | 25 | 89 |
| Glyma.02G245700 | Calmodulin and related proteins (EF-Hand superfamily) | Signal Transduction | 131 | 711 |
| Glyma.04G194800 | Calmodulin and related proteins (EF-Hand superfamily) | Signal Transduction | 221 | 1264 |
| Glyma.02G002100 | Calmodulin and related proteins (EF-Hand superfamily) | Signal Transduction | 188 | 1266 |
| Glyma.19G244300 | Calmodulin and related proteins (EF-Hand superfamily) | Signal Transduction | 191 | 2286 |
| Glyma.16G142500 | Calmodulin and related proteins (EF-Hand superfamily) | Signal Transduction | 229 | 3426 |
| Glyma.15G212400 | Cell cycle control protein | Signal Transduction | 119 | 786 |
| Glyma.02G257600 | Dual-specificity tyrosine-phosphorylation regulated kinase | Signal Transduction | 277 | 775 |
| Glyma.18G298300 | Multifunctional chaperone (14-3-3 family) | Signal Transduction | 372 | 5003 |
| Glyma.17G153800 | Serine/threonine phosphatase | Signal Transduction | 161 | 2794 |
| Glyma.12G140200 | Serine/threonine protein kinase | Signal Transduction | 515 | 15 |
| Glyma.09G014900 | Serine/threonine protein kinase | Signal Transduction | 38 | 191 |
| Glyma.20G138500 | Serine/threonine protein kinase | Signal Transduction | 42 | 354 |
| Glyma.13G073900 | Serine/threonine protein kinase | Signal Transduction | 364 | 411 |
| Glyma.18G214000 | Serine/threonine protein kinase | Signal Transduction | 409 | 939 |
| Glyma.18G275700 | Serine/threonine protein kinase | Signal Transduction | 381 | 2800 |
| Glyma.18G216800 | Serine/threonine protein kinase | Signal Transduction | 367 | 136 |
| Glyma.04G230500 | Serine/threonine protein kinase | Signal Transduction | 459 | 150 |
| Glyma.10G109200 | Serine/threonine protein kinase | Signal Transduction | 304 | 331 |
| Glyma.10G126700 | Serine/threonine protein kinase | Signal Transduction | 449 | 408 |
| Glyma.20G140100 | Serine/threonine protein kinase | Signal Transduction | 219 | 417 |
| Glyma.04G042400 | Serine/threonine protein kinase | Signal Transduction | 150 | 448 |
| Glyma.20G054500 | Serine/threonine protein kinase | Signal Transduction | 307 | 753 |
| Glyma.08G235900 | Serine/threonine protein kinase | Signal Transduction | 368 | 1434 |
| Glyma.09G089700 | Serine/threonine protein kinase | Signal Transduction | 322 | 1826 |
| Glyma.20G139700 | Serine/threonine protein kinase | Signal Transduction | 441 | 4831 |
| Glyma.17G056900 | Serine/threonine protein kinase | Signal Transduction | 355 | 211 |
| Glyma.20G138800 | Serine/threonine protein kinase | Signal Transduction | 374 | 932 |
| Glyma.13G102200 | Serine/threonine protein kinase | Signal Transduction | 284 | 1459 |
| Glyma.05G237100 | Serine/threonine protein kinase | Signal Transduction | 296 | 2831 |
| Glyma.02G009100 | Serine/threonine protein phosphatase | Signal Transduction | 511 | 3557 |
| Glyma.12G170100 | SOK1 kinase belonging to the STE20/SPS1/GC kinase family | Signal Transduction | 408 | 1017 |
| Glyma.05G036600 | Tyrosine kinase specific for activated (GTP-bound) p21cdc42Hs | Signal Transduction | 590 | 2130 |
| Glyma.07G142000 | Uncharacterized conserved protein | Signal Transduction | 521 | 3604 |
| Glyma.14G016300 | CCCH-type Zn-finger protein | Transcription | 431 | 1274 |
| Glyma.03G060200 | HMG box-containing protein | Transcription | 583 | 1173 |
| Glyma.15G272400 | Predicted E3 ubiquitin ligase | Transcription | 519 | 4432 |
| Glyma.11G246100 | Predicted E3 ubiquitin ligase | Transcription | 407 | 1380 |
| Glyma.03G179300 | Predicted E3 ubiquitin ligase | Transcription | 331 | 3510 |
| Glyma.05G178200 | Uncharacterized conserved protein, contains IPT/TIG domain | Transcription | 477 | 3055 |
| Glyma.16G062600 | Amino acid transporters | Transporter | 109 | 140 |
| Glyma.19G076800 | Amino acid transporters | Transporter | 76 | 298 |
| Glyma.05G043100 | Amino acid transporters | Transporter | 315 | 4608 |
| Glyma.10G132300 | Ammonia permease | Transporter | 72 | 223 |
| Glyma.05G211600 | Copper chaperone | Transporter | 311 | 241 |
| Glyma.07G065800 | Copper chaperone | Transporter | 579 | 733 |
| Glyma.06G191300 | Lipid exporter ABCA1 and related proteins, ABC superfamily | Transporter | 114 | 312 |
| Glyma.18G050400 | Mitochondrial Fe2 transporter MMT1 and related transporters (cation diffusion facilitator superfamily) | Transporter | 97 | 1023 |
| Glyma.10G276700 | Monocarboxylate transporter | Transporter | 158 | 109 |
| Glyma.20G112900 | Monocarboxylate transporter | Transporter | 604 | 798 |
| Glyma.10G019000 | Multidrug resistance-associated protein/mitoxantrone resistance protein, ABC superfamily | Transporter | 286 | 101 |
| Glyma.02G008000 | Multidrug/pheromone exporter, ABC superfamily | Transporter | 446 | 558 |
| Glyma.09G056300 | Plasma membrane H<sup>+</sup>-transporting ATPase | Transporter | 212 | 184 |
| Glyma.13G097900 | Plasma membrane H<sup>+</sup>-transporting ATPase | Transporter | 167 | 317 |
| Glyma.14G134100 | Plasma membrane H<sup>+</sup>-transporting ATPase | Transporter | 423 | 788 |
| Glyma.15G011900 | Pleiotropic drug resistance proteins (PDR1-15), ABC superfamily | Transporter | 233 | 12 |
| Glyma.13G361900 | Pleiotropic drug resistance proteins (PDR1-15), ABC superfamily | Transporter | 26 | 79 |
| Glyma.01G223600 | Predicted K<sup>+</sup> antiporter | Transporter | 332 | 1386 |
| Glyma.13G131200 | Predicted K<sup>+</sup> antiporter | Transporter | 269 | 1889 |
| Glyma.01G156200 | Predicted membrane protein | Transporter | 546 | 283 |
| Glyma.01G238800 | Predicted transporter (major facilitator superfamily) | Transporter | 128 | 69 |
| Glyma.08G035300 | Predicted transporter (major facilitator superfamily) | Transporter | 518 | 126 |
| Glyma.19G020000 | Prohibitins and stomatins of the PID superfamily | Transporter | 283 | 291 |
| Glyma.13G065000 | Prohibitins and stomatins of the PID superfamily | Transporter | 171 | 1582 |
| Glyma.02G021400 | Prohibitins and stomatins of the PID superfamily | Transporter | 406 | 2729 |
| Glyma.02G129500 | P-type ATPase | Transporter | 241 | 925 |

**Additional file 1: Table S1:** List of the soybean genes up regulated 10 days after *P. pachyrhizi* inoculation. Soybean genes were identified in Tremblay *et al.*, 2010 and Tremblay *et al.*, 2011. Genes were re-annotated with the last soybean genome notation available: Glycine max 275 William 82 (from https://genome.jgi.doe.gov/portal/soybean/soybean.home.html). The genes were ranked according to their relative expression level compared to the mock inoculation.

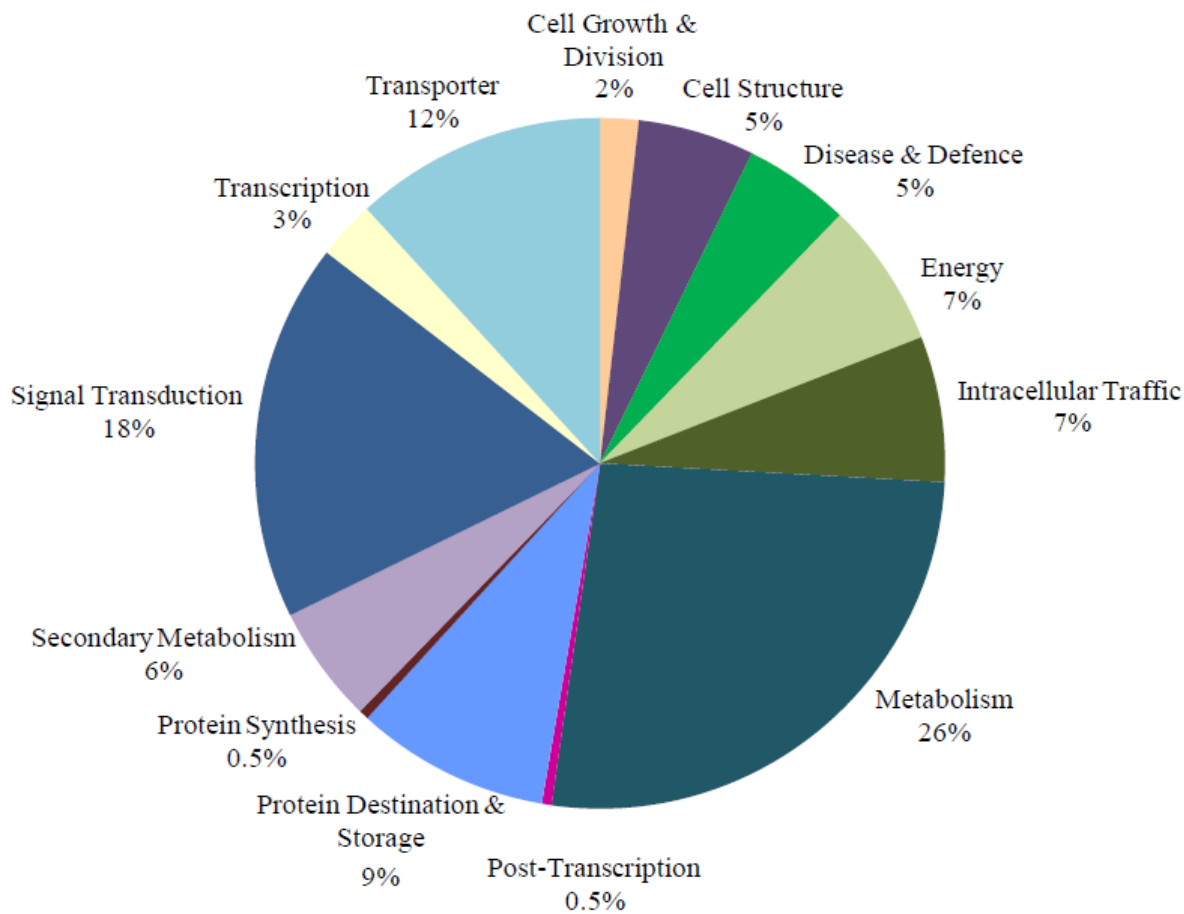

**Additional file 2: Figure S1:** Functional categories of the 220 soybean genes up-regulated 10 days post inoculation with *P. pachyrhizi* and identified in Tremblay *et al.*, 2010 and Tremblay *et al.*, 2011.

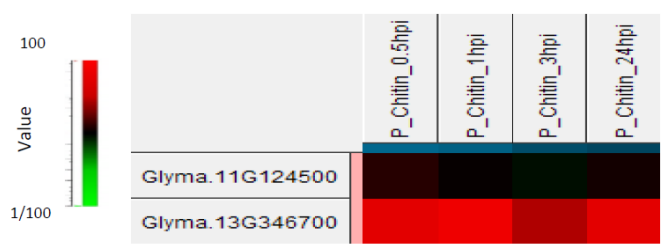

**Additional file 3: Figure S2:** Relative expression of *Glyma.13G346700* and *Glyma.11G124500* compared to control plants (untreated) at 0.5, 1, 3, and 24 hours post chitinheptaose (DP7) treatment. Black boxes represent no change in genes expression compared with the control plants, red boxes indicate upregulation by DP7 treatment. The plants were sprayed with 200 ppm of DP7 or water until run-off. They were incubated in growth chamber temperature (24°C, 16 h light/8 h night, light intensity 15 $\mu\text{E}\cdot\text{m}^{-2}\cdot\text{s}^{-1}$ and 80% relative humidity) for 24 hours.

| Proteins homolog | Predicted function | % of identity |
| --- | --- | --- |
| *Glycine soja* KHN06178.1 | Endochitinase PR4 | 100 |
| *Cicer arietium* XP_012570981.1 | Chitinase | 85 |
| *Vigna angularis* BAT90863.1 | Hypothetical protein | 82 |
| *Lotus japonicus* AFK37127.1 | Unknown | 80 |
| *Phaseolus vulgaris* XP_007131917.1 | Hypothetical protein | 80 |
| *Vigna radiata* XP_022642874.1 | Uncharacterized protein | 78 |
| *Arachis hypogaea* EP3 XP025686362 | Endochitinase | 75 |
| *Arachis ipaensis* XP_016187096.1 | Endochitinase EP3 | 75 |
| *Arachis duranensis* XP015952096 | Endochitinase EP3 | 75 |
| *Cajanus cajan* XP_020224332.1 | Endochitinase PR4-like | 69 |
| *Medicago sativa* ACL36992.1 | Chitinase class IV | 66 |
| *Medicago truncatula* XP_003597548.2 | Endochitinase PR4 | 66 |
| *Trifolium pratense* PNY03482.1 | Endochitinase PR4-like | 64 |

**Additional file 4: Figure S3:** GmCHIT protein KRH29572.1 homologs, their functions and % of identity. From *BLASTP* analyze (NCBI).

1 XP\_0071319 -----mskVAGFvTlFMvNMNVPKtaSA1-----
2 BAT90863.1 ---MlqvantakVAGFvTlFMvNMNVPKtVSAa---
3 XP\_0226428 ---MlqvantakVAGFvTlFMvNMNVPKtVSAa---
4 XP\_0256863 ---M-MakRLl1-----MgiwvMdg---MVk-----anG---
5 XP\_0161870 ---M-MakRLl1-----MgiwvMdg---MVk-----anG---
6 XP\_0159520 ---M-MakRLl1-----MgiwvMdg---MVk-----anG---
7 XP\_0194143 ---M-MakRLl1-----MgiwvMdg---MVk-----anG---
8 PNY03482.1 ---M-MakRLl1-----MgiwvMdg---MVk-----anG---
9 ACL36992.1 ---M-MakRLl1-----MgiwvMdg---MVk-----anG---
10 XP\_0035975 ---M-MakRLl1-----MgiwvMdg---MVk-----anG---
11 AFR37127.1 ---M-MakRLl1-----MgiwvMdg---MVk-----anG---
12 XP\_0125709 ---M-MakRLl1-----MgiwvMdg---MVk-----anG---
13 XP\_0202243 ---M-MakRLl1-----MgiwvMdg---MVk-----anG---
14 KRH29572.1 ---M-MakRLl1-----MgiwvMdg---MVk-----anG---
15 KHM06178.1 ---M-MakRLl1-----MgiwvMdg---MVk-----anG---
16 XP\_0071319 -----hiADIVT-qFFNnIIInkADgCpGKaFYSRDAFLKa
17 BAT90863.1 -----Nid-IVTqGFFdmIIInQADgCpGKNFYSRDAFLKa
18 XP\_0226428 -----hidkIVTqGFFdmIIInQADgCpGKNFYSRDAFLKa
19 XP\_0256863 -----sVgDIVTqGFFNnIIInkADgCpGKaFYSRDAFLKa
20 XP\_0161870 -----sVgDIVTqGFFNnIIInkADgCpGKaFYSRDAFLKa
21 XP\_0159520 -----sVgDIVTqGFFNnIIInkADgCpGKaFYSRDAFLKa
22 XP\_0194143 -----sVgDIVTqGFFNnIIInkADgCpGKaFYSRDAFLKa
23 PNY03482.1 -----sVgDIVTqGFFNnIIInkADgCpGKaFYSRDAFLKa
24 ACL36992.1 -----sVgDIVTqGFFNnIIInkADgCpGKaFYSRDAFLKa
25 XP\_0035975 -----sVgDIVTqGFFNnIIInkADgCpGKaFYSRDAFLKa
26 AFR37127.1 -----sVgDIVTqGFFNnIIInkADgCpGKaFYSRDAFLKa
27 XP\_0125709 -----sVgDIVTqGFFNnIIInkADgCpGKaFYSRDAFLKa
28 XP\_0202243 -----sVgDIVTqGFFNnIIInkADgCpGKaFYSRDAFLKa
29 KRH29572.1 -----sVgDIVTqGFFNnIIInkADgCpGKaFYSRDAFLKa
30 KHM06178.1 -----sVgDIVTqGFFNnIIInkADgCpGKaFYSRDAFLKa
31 XP\_0071319 hNSYp-FCaL-naDDSKRE-AAAFAHFTHEsGHFCYIEEIGAS-DYCDENTEYPCaP
32 BAT90863.1 hNSYp-FCaL-naDDSKRE-AAAFAHFTHEsGHFCYIEEIGAS-DYCDENTEYPCaP
33 XP\_0226428 hNSYp-FCaL-naDDSKRE-AAAFAHFTHEsGHFCYIEEIGAS-DYCDENTEYPCaP
34 XP\_0256863 hNSYp-FCaL-naDDSKRE-AAAFAHFTHEsGHFCYIEEIGAS-DYCDENTEYPCaP
35 XP\_0161870 hNSYp-FCaL-naDDSKRE-AAAFAHFTHEsGHFCYIEEIGAS-DYCDENTEYPCaP
36 XP\_0159520 hNSYp-FCaL-naDDSKRE-AAAFAHFTHEsGHFCYIEEIGAS-DYCDENTEYPCaP
37 XP\_0194143 hNSYp-FCaL-naDDSKRE-AAAFAHFTHEsGHFCYIEEIGAS-DYCDENTEYPCaP
38 PNY03482.1 hNSYp-FCaL-naDDSKRE-AAAFAHFTHEsGHFCYIEEIGAS-DYCDENTEYPCaP
39 ACL36992.1 hNSYp-FCaL-naDDSKRE-AAAFAHFTHEsGHFCYIEEIGAS-DYCDENTEYPCaP
40 XP\_0035975 hNSYp-FCaL-naDDSKRE-AAAFAHFTHEsGHFCYIEEIGAS-DYCDENTEYPCaP
41 AFR37127.1 hNSYp-FCaL-naDDSKRE-AAAFAHFTHEsGHFCYIEEIGAS-DYCDENTEYPCaP
42 XP\_0125709 hNSYp-FCaL-naDDSKRE-AAAFAHFTHEsGHFCYIEEIGAS-DYCDENTEYPCaP
43 XP\_0202243 hNSYp-FCaL-naDDSKRE-AAAFAHFTHEsGHFCYIEEIGAS-DYCDENTEYPCaP
44 KRH29572.1 hNSYp-FCaL-naDDSKRE-AAAFAHFTHEsGHFCYIEEIGAS-DYCDENTEYPCaP
45 KHM06178.1 hNSYp-FCaL-naDDSKRE-AAAFAHFTHEsGHFCYIEEIGAS-DYCDENTEYPCaP
46 XP\_0071319 KGYGRCPIQLSWNFNYGPAGqNIGFDGLNAPETVANDPVVSFKTALWYWMNVVRPVINQ
47 BAT90863.1 KGYGRCPIQLSWNFNYGPAGqNIGFDGLNAPETVANDPVVSFKTALWYWMNVVRPVINQ
48 XP\_0226428 KGYGRCPIQLSWNFNYGPAGqNIGFDGLNAPETVANDPVVSFKTALWYWMNVVRPVINQ
49 XP\_0256863 KGYGRCPIQLSWNFNYGPAGqNIGFDGLNAPETVANDPVVSFKTALWYWMNVVRPVINQ
50 XP\_0161870 KGYGRCPIQLSWNFNYGPAGqNIGFDGLNAPETVANDPVVSFKTALWYWMNVVRPVINQ
51 XP\_0159520 KGYGRCPIQLSWNFNYGPAGqNIGFDGLNAPETVANDPVVSFKTALWYWMNVVRPVINQ
52 XP\_0194143 KGYGRCPIQLSWNFNYGPAGqNIGFDGLNAPETVANDPVVSFKTALWYWMNVVRPVINQ
53 PNY03482.1 KGYGRCPIQLSWNFNYGPAGqNIGFDGLNAPETVANDPVVSFKTALWYWMNVVRPVINQ
54 ACL36992.1 KGYGRCPIQLSWNFNYGPAGqNIGFDGLNAPETVANDPVVSFKTALWYWMNVVRPVINQ
55 XP\_0035975 KGYGRCPIQLSWNFNYGPAGqNIGFDGLNAPETVANDPVVSFKTALWYWMNVVRPVINQ
56 AFR37127.1 KGYGRCPIQLSWNFNYGPAGqNIGFDGLNAPETVANDPVVSFKTALWYWMNVVRPVINQ
57 XP\_0125709 KGYGRCPIQLSWNFNYGPAGqNIGFDGLNAPETVANDPVVSFKTALWYWMNVVRPVINQ
58 XP\_0202243 KGYGRCPIQLSWNFNYGPAGqNIGFDGLNAPETVANDPVVSFKTALWYWMNVVRPVINQ
59 KRH29572.1 KGYGRCPIQLSWNFNYGPAGqNIGFDGLNAPETVANDPVVSFKTALWYWMNVVRPVINQ
60 KHM06178.1 KGYGRCPIQLSWNFNYGPAGqNIGFDGLNAPETVANDPVVSFKTALWYWMNVVRPVINQ
61 XP\_0071319 GFGATIRAINQ@LECDqgNPaTVQARVNHYYTQYCSQLGVAPGDNLTc-----
62 BAT90863.1 GFGATIRAINQ@LECDqgNPaTVQARVNHYYTQYCSQLGVAPGDNLTc-----
63 XP\_0226428 GFGATIRAINQ@LECDqgNPaTVQARVNHYYTQYCSQLGVAPGDNLTc-----
64 XP\_0256863 GFGATIRAINQ@LECDqgNPaTVQARVNHYYTQYCSQLGVAPGDNLTc-----
65 XP\_0161870 GFGATIRAINQ@LECDqgNPaTVQARVNHYYTQYCSQLGVAPGDNLTc-----

```
66 XP_0159520 GFGATIRAINGQLECDGANPTTVQARVNYXQYCSQLGVdPGDMLTC-----
67 XP_0194143 GFGATIRAINGaLECDGgNPpTVQARVNYTQYCSQLGVatGDMLTC-----
68 PMY03482.1 GFGATIRAINGkLECDGANPTTVQARVdYYkQYCSQLGVaPGDMLTC-----
69 ACL36992.1 GFGATIRAING-LECDGANPSTVQtrVgYYTQYCSaLGVAfGDMLTC-----
70 XP_0035975 GFGATIRAING-LECDGANPSTVQtrVgYYTQYCSaLGVAfGDMLTC-----
71 AFK37127.1 GFGATIRAINGQLECDGANaTVQARVNYTQYCSQLGVaPGDMLT-----
72 XP_0125709 GFGATIRAINGQLECDGANPpTVQARVvYYTQYCSQLGVaPGDMLTC-----
73 XP_0202243 GFGATIRAINGQLECDGANPTTVQARVNYTQYCSQLGVatGDMLTC-----
74 KRH29572.1 GFGATIRAING-LECDGANPSTVQARVNYTQYCSQLGVaPGDMLTC-----
75 KHM06178.1 GFGATIRAING-LECDGANPSTVQARVNYTQYCSQLGVaPGDMLTC-----

76 XP_0071319 -----
77 BAT90863.1 -----
78 XP_0226428 gacvfivitqfd=ilhqanwnkkvqnmntgvt=vwvafialmligcqqgacygtdeanh
79 XP_0256863 v-----
80 XP_0161870 -----
81 XP_0159520 -----
82 XP_0194143 -----
83 PMY03482.1 -----
84 ACL36992.1 -----
85 XP_0035975 -----
86 AFK37127.1 -----
87 XP_0125709 -----
88 XP_0202243 -----
89 KRH29572.1 -----
90 KHM06178.1 -----

91 XP_0071319 -----
92 BAT90863.1 -----
93 XP_0226428 avadivtteffnnifdagdddadcpqknfy=qafihalnaykqfg=agavddak=aiiaa
94 XP_0256863 -----
95 XP_0161870 -----
96 XP_0159520 -----
97 XP_0194143 -----
98 PMY03482.1 -----
99 ACL36992.1 -----
100 XP_0035975 -----
101 AFK37127.1 -----
102 XP_0125709 -----
103 XP_0202243 -----
104 KRH29572.1 -----
105 KHM06178.1 -----

106 XP_0071319 -----
107 BAT90863.1 -----
108 XP_0226428 afahftthetghfcyieesegaskdycdaskadapcaadkayyg=gpiqlkwnynygaa
109 XP_0256863 g-----
110 XP_0161870 -----
111 XP_0159520 -----
112 XP_0194143 -----
113 PMY03482.1 -----
114 ACL36992.1 -----
115 XP_0035975 -----
116 AFK37127.1 -----
117 XP_0125709 -----
118 XP_0202243 -----
119 KRH29572.1 -----
120 KHM06178.1 -----

121 XP_0071319 -----
122 BAT90863.1 -----
123 XP_0226428 esigfdgllkapetvgadpvrvaftalwy=tenvapvmaqgfgeti=amkgeveadgggnpd
124 XP_0256863 -----
125 XP_0161870 -----
126 XP_0159520 -----
127 XP_0194143 -----
128 PMY03482.1 -----
129 ACL36992.1 -----
130 XP_0035975 -----
131 AFK37127.1 -----
```

|  |  |  |
| --- | --- | --- |
| 132 | XP_0125709 | ----- |
| 133 | XP_0202243 | ----- |
| 134 | KRH29572.1 | ----- |
| 135 | KHM06178.1 | ----- |
| 136 | XP_0071319 | ===== |
| 137 | BAT90863.1 | ===== |
| 138 | XP_0226428 | avqaevdyrtqycaqlgwapgdnlta |
| 139 | XP_0236863 | ===== |
| 140 | XP_0161870 | ===== |
| 141 | XP_0159320 | ===== |
| 142 | XP_0194143 | ===== |
| 143 | PMY03482.1 | ===== |
| 144 | ACL36992.1 | ===== |
| 145 | XP_0035975 | ===== |
| 146 | AFK37127.1 | ===== |
| 147 | XP_0125709 | ===== |
| 148 | XP_0202243 | ===== |
| 149 | KRH29572.1 | ----- |
| 150 | KHM06178.1 | ===== |

26

27 **Additional file 5: Figure S4:** Multiple Sequence Alignment of GmCHIT (**KRH29572.1**) with plant
28 homologous proteins (from *BLASTP* analysis). Similar residues are colored according to BLOSUM62
29 score: Max: 3.0 Low: 0.5.

30

a

| Factor or Site Name | Location | strain | Signal Sequence | description |
| --- | --- | --- | --- | --- |
| ARFAT | 85 | (+) | TGTCTC | AUX |
| AUXREPSIAA4 | 3278 | (-) | KGTCCCAT | AUX |
| CAREOSREP1 | 19 | (+) | CAACTC | GA |
| CAREOSREP1 | 2019 | (-) | CAACTC | GA |
| GT1GMSCAM4 | 225 | (-) | GAAAAA | pathogen salt induction |
| GT1GMSCAM4 | 566 | (+) | GAAAAA | pathogen salt induction |
| GT1GMSCAM4 | 1644 | (+) | GAAAAA | pathogen salt induction |
| GT1GMSCAM4 | 1922 | (-) | GAAAAA | pathogen salt induction |
| GT1GMSCAM4 | 2331 | (+) | GAAAAA | pathogen salt induction |
| GT1GMSCAM4 | 2533 | (+) | GAAAAA | pathogen salt induction |
| GT1GMSCAM4 | 2769 | (-) | GAAAAA | pathogen salt induction |
| GT1GMSCAM4 | 2860 | (-) | GAAAAA | pathogen salt induction |
| GT1GMSCAM4 | 2878 | (-) | GAAAAA | pathogen salt induction |
| GT1GMSCAM4 | 3031 | (-) | GAAAAA | pathogen salt induction |
| MYBST1 | 122 | (-) | GGATA | MYB TF (biotic/abiotic response) |
| MYBST1 | 798 | (-) | GGATA | MYB TF (biotic/abiotic response) |
| MYBST1 | 1131 | (-) | GGATA | MYB TF (biotic/abiotic response) |
| MYBST1 | 1819 | (-) | GGATA | MYB TF (biotic/abiotic response) |
| MYBST1 | 1888 | (-) | GGATA | MYB TF (biotic/abiotic response) |
| TATABOX2 | 332 | (+) | TATAAAT | TATA box |
| WBOXPCWRKY1 | 485 | (+) | TTTGACY | W_box |
| WBOXPCWRKY1 | 517 | (+) | TTTGACY | W_box |
| WBOXPCWRKY1 | 1276 | (+) | TTTGACY | W_box |
| WBOXPCWRKY1 | 3102 | (-) | TTTGACY | W_box |
| WBOXPCWRKY1 | 3233 | (-) | TTTGACY | W_box |
| WBOXATNPR1 | 401 | (-) | TTGAC | W_box |
| WBOXATNPR1 | 410 | (+) | TTGAC | W_box |
| WBOXATNPR1 | 883 | (-) | TTGAC | W_box |
| WBOXATNPR1 | 1672 | (-) | TTGAC | W_box |
| WBOXHVISO1 | 117 | (+) | TGACT | W_box |
| WBOXHVISO1 | 479 | (-) | TGACT | W_box |
| WBOXHVISO1 | 504 | (+) | TGACT | W_box |
| WBOXHVISO1 | 696 | (+) | TGACT | W_box |
| WBOXHVISO1 | 1354 | (+) | TGACT | W_box |
| WBOXHVISO1 | 3169 | (-) | TGACT | W_box |
| WBOXNTERF3 | 1380 | (-) | TGACY | W_box |
| WRKY71OS | 480 | (-) | TGAC | W_box |
| WRKY71OS | 1175 | (-) | TGAC | W_box |
| WRKY71OS | 1589 | (-) | TGAC | W_box |
| WRKY71OS | 1802 | (-) | TGAC | W_box |
| WRKY71OS | 2385 | (+) | TGAC | W_box |

b

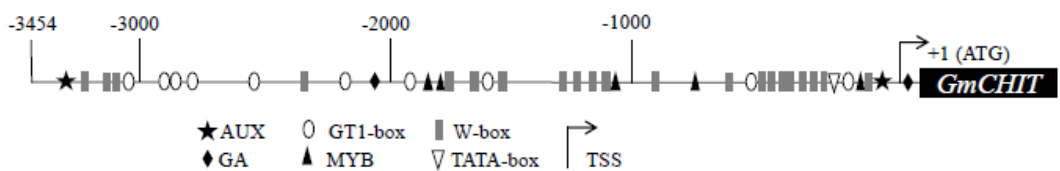

**Additional file 6: Figure S5:** The *GmCHIT1* promoter and potential *cis*-regulatory elements identified. (a) List of the *cis*-regulatory elements related to pathogen infection identified in the *GmCHIT1* promoter. (b) Map of the *GmCHIT1* promoter. AUX: auxin responsive element, GA: gibberellic acid responsive elements, MYB: MYB recognition elements, GT1-box: pathogen and NaCl responsive elements, W-box: pathogen responsive elements, TSS: transcription start site.

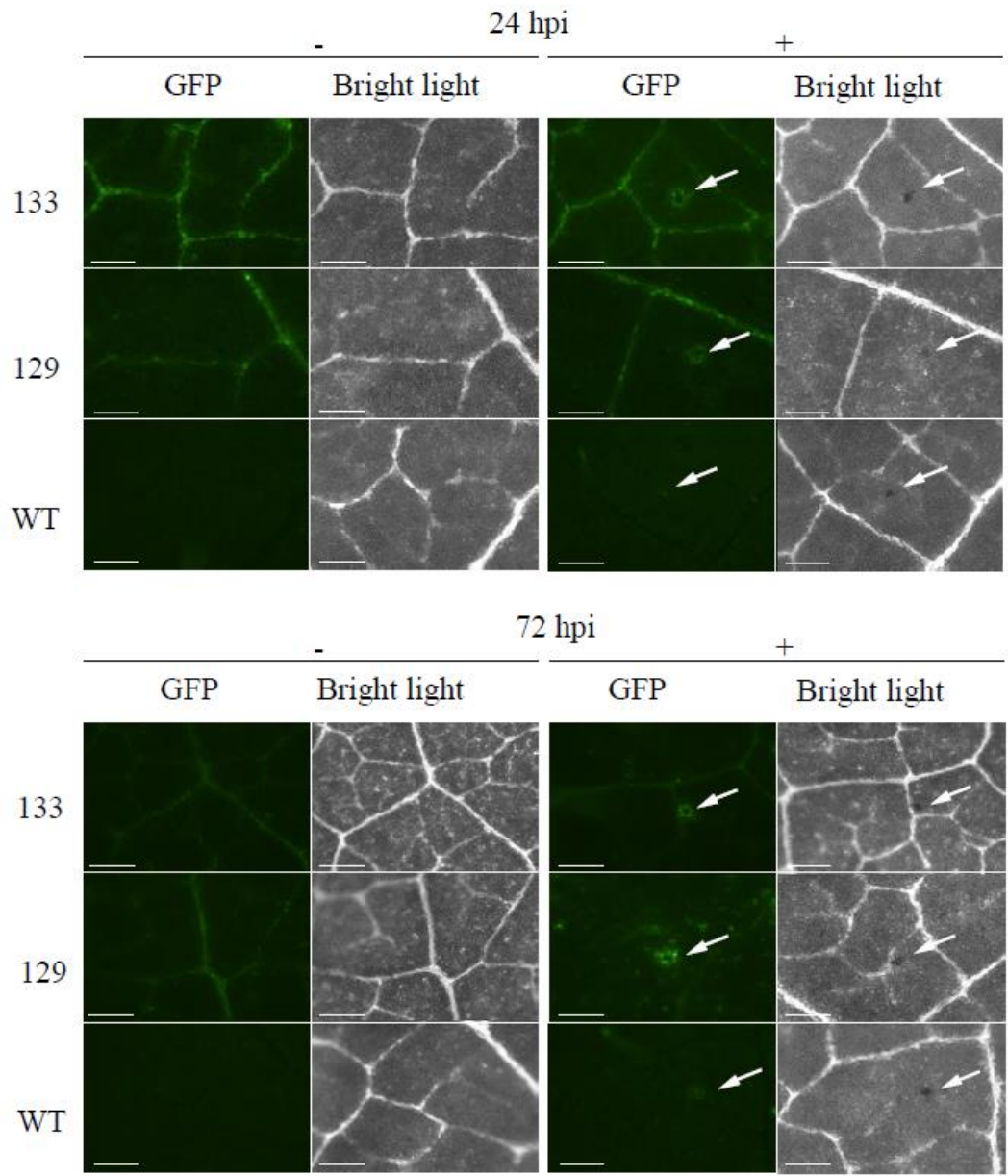

**Additional file 7: Figure S6:** Detection of GFP fluorescence in stable transgenic soybeans. Leaves of two T1 lines (129 and 133) transformed with the p*GmCHIT1*:GFP construction were observed using a dissection scope (Leica Z16 APO) under GFP filter and bright light at 24 and 72 hours after. *P. pachyrhizi* inoculation (+) or mock treatment (-). Arrows indicate the inoculation spots. Bar-scales represent 200 μm.

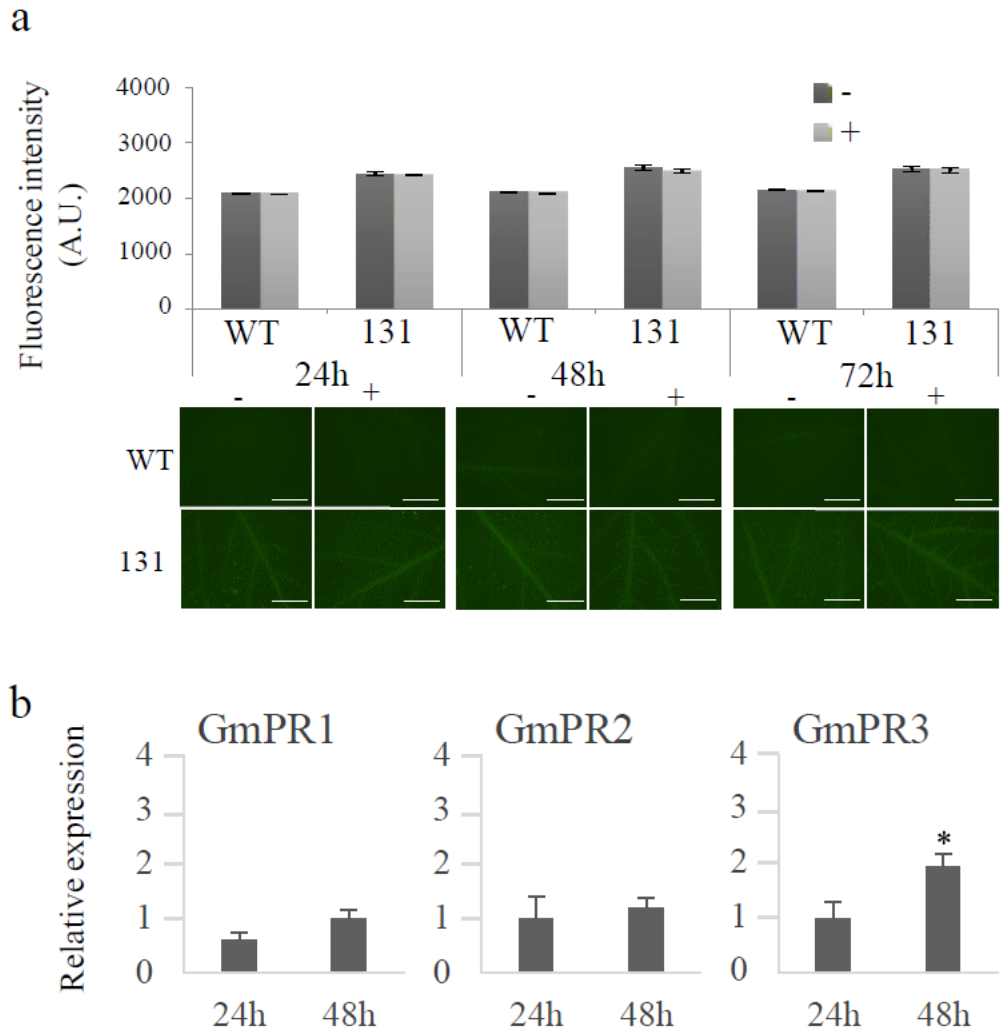

**Additional file 8: Figure S7: *GmCHIT1* promoter expression following salicylic acid treatment.** (a) GFP fluorescence in 131 line (p*GmCHIT1*:GFP) and WT detached leaves following SA (+) or mock (-) treatments. Graphics represent the fluorescence intensity measured with MetaMorph software *via* grayscale value. Mean of 20 biological replicates ± standard errors. No significant difference between treated (+) and untreated (-) leaves (Student's t-test, $p < 0.05$). Representative images of the observed fluorescence are shown under the graphs. Bar-scales represent 5mm. Observations were realized at 24, 48 and 72 hours after hormonal treatment with a dissection scope (Leica Z16 APO) under GFP filter. (b) Relative expression of *GmPR1* (GenBank: BU5773813), *GmPR2* (GenBank: M37753), *GmPR3* (GenBank: AF202731) in event 131 (p*GmCHIT1*:GFP) after SA treatment. Transcript accumulation at 24 and 48 hours compared to that in the mock-treated plants. The actin (GenBank: NM\_001289231.2) and an elongation factor (GenBank: NM\_001249608.2) encoding genes were used as references <sup>68</sup>.

Three independent biological replicates ± standard deviations. \*: significant difference between treated and untreated leaves determined by a Student's t-test ($p < 0.05$).

| Amplified sequence | Primer name | Primer sequence | Amplicon length | Primer use |
| --- | --- | --- | --- | --- |
| GFP | gfp F | GAATATTCACGTGAATTCATGGTGAGCAAGGGCGAGGAGC | 759bp | PCR for cloning |
|  | gfp R | ACCTGCAGGCCACAGCTGTGGTTACTTGTACAGCTCGTCCATGCCG |  |  |
| PDF1.2 promoter | pdf1.2_F | GTAATACGCGTCTTAAGACGTCTGGGACAAGCTTTATATGCAG | 1253bp |  |
|  | pdf1.2_R | GGACGTAACATGAATTCGATGATTATTACTATTTTG |  |  |
| Actin | GmACTIN_F | CGGTGGTTCTATCTTGGCATC | 142bp | quantitative PCR |
|  | GmACTIN_R | GTCTTTCGCTTCAATAACCCTA |  |  |
| Hypothetical protein | GmUKN2_F* | GCCTGGATACCTGCTCAAG | 79bp |  |
|  | GmUKN2_R* | ACCTCCTCCTCAAACCTCTCTG |  |  |
| Chitinase | GmCHIT_F | GAGATTAAACGGTGCATCAGG | 330bp |  |
|  | GmCHIT_R | ATTAACACGAGCCTGAACAGTACT |  |  |
| GmPR1 | GmPR1_F** | AACTATGCTCCCCCTGGCAACTATATTG | 76bp |  |
|  | GmPR1_R** | TCTGAAGTGGTAGCTTCTACATCGAAACAA |  |  |
| GmPR2 | GmPR2_F*** | GTCTCCTTCGGTGGTAGTG | 104bp |  |
|  | GmPR2_R*** | ACCCTCCTCCTGCTTTCTC |  |  |
| GmPR3 | GmPR3_F*** | GCATTGGTCTGGATTTG | 115bp |  |
|  | GmPR3_R*** | GGCTTGATGGCTTGTTTC |  |  |
| Elongation factor | GmELF_F | GTTGAAAAGCCAGGGGACA | 79bp |  |
|  | GmELF_R | TCTTACCCCTTGAGCGTGG |  |  |

**Additional file 9: Table S2:** Primers used for PCR and qPCR. \*from Hirschburger et al., 2015<sup>68</sup>, \*\* from Mazarei et al., 2007<sup>62</sup>, \*\*\* from Zhong et al., 2014<sup>69</sup>.
